# Benchmarking the accuracy of structure-based binding affinity predictors on Spike-ACE2 Deep Mutational Interaction Set

**DOI:** 10.1101/2022.04.18.488633

**Authors:** Burcu Ozden, Eda Şamiloğlu, Atakan Özsan, Mehmet Erguven, Mehdi Koşaca, Melis Oktayoğlu, Can Yükrük, Nazmiye Arslan, Gökhan Karakülah, Ayşe Berçin Barlas, Büşra Savaş, Ezgi Karaca

## Abstract

Since the start of COVID-19 pandemic, a huge effort has been devoted to understanding the Spike (SARS-CoV-2)-ACE2 recognition mechanism. To this end, two deep mutational scanning studies traced the impact of all possible mutations across Receptor Binding Domain (RBD) of Spike and catalytic domain of human ACE2. By concentrating on the interface mutations of these experimental data, we benchmarked six commonly used structure-based binding affinity predictors (FoldX, EvoEF1, MutaBind2, SSIPe, HADDOCK, and UEP). These predictors were selected based on their user-friendliness, accessibility, and speed. As a result of our benchmarking efforts, we observed that none of the methods could generate a meaningful correlation with the experimental binding data. The best correlation is achieved by FoldX (R = -0.51). Also, when we simplified the prediction problem to a binary classification, i.e., whether a mutation is enriching or depleting the binding, we showed that the highest accuracy is achieved by FoldX with 64% success rate. Surprisingly, on this set, simple energetic scoring functions performed significantly better than the ones using extra evolutionary-based terms, as in Mutabind and SSIPe. Furthermore, we also demonstrated that recent AI approaches, mmCSM-PPI and TopNetTree, yielded comparable performances to the force field-based techniques. These observations suggest plenty of room to improve the binding affinity predictors in guessing the variant-induced binding profile changes of a host-pathogen system, such as Spike-ACE2. To aid such improvements we provide our benchmarking data at https://github.com/CSB-KaracaLab/RBD-ACE2-MutBench with the option to visualize our mutant models at https://rbd-ace2-mutbench.github.io/

## INTRODUCTION

At the beginning of 21^st^ century, the emergence of Severe Acute Respiratory Syndrome Coronavirus (SARS-CoV)[1] and Middle East Respiratory Syndrome Coronavirus[2] led to serious public health concerns. Evolving from these viruses, during late 2019, a new SARS virus, SARS-CoV-2, caused the most severe pandemic of the 21^st^ century[3]. SARS-CoV- 2 infection is initiated upon having its Spike protein interacting with the host Angiotensin Converting 2 (ACE2) enzyme[4]. The widespread infection of SARS-CoV-2 compared to its predecessors was linked to higher binding affinity of Spike to ACE2[5]. Relatedly, alpha, beta, gamma, eta, iota, kappa, lambda, mu, and omicron SARS-CoV-2 variants were shown to have at least one mutation across the Spike-ACE2 interface[6]. This realization placed the characterization of interfacial Spike-ACE2 mutations at the center of COVID-19-related research. Within this context, in 2020, two deep mutational scanning (DMS) studies explored how Spike/ACE2 variants impact Spike-ACE2 interactions[7,8]. In these DMS studies, the residues on the Receptor Binding Domain (RBD) of Spike and the catalytic domain of human ACE2 were mutated into other 19 amino acid possibilities, followed by tracing of new RBD-ACE2 binding profiles.

In parallel to these experimental efforts, a handful of structure-based computational studies employed a comprehensive investigation of variation across the RBD-ACE2 interface (Table 1). Three such studies utilized two fast and user-friendly tools, FoldX and HADDOCK [9–11]. Blanco et al. used FoldX with the inclusion of water molecules (FoldXwater) to trace the binding enhancing RBD and ACE2 mutations [9]. From their 21 binding enhancing mutation predictions, nine of them were confirmed as affinity enhancing by the DMS set. Rodrigues et al. investigated the impact of ACE2 orthologs on their RBD binding with HADDOCK, where they proposed five significantly affinity improving ACE2 mutations[10]. Among these, D30E and A387E were shown to be affinity enhancing in the ACE2 DMS set as well. Complementary to this study, Sorokina et al. performed computational alanine scanning on ACE2 with HADDOCK[11]. Here, three out of five mutations (N49A, R393A, and P389A) were classified as binding enriching both by the computational predictions and the DMS set. Two other studies made use of elaborate simulation techniques, mainly molecular dynamics simulations. Laurini et al. performed molecular mechanics/Poisson−Boltzmann alanine scanning and molecular dynamics simulations to find structurally and energetically critical RBD-ACE2 residues[12]. They proposed eight hotspot positions on RBD and ACE2, where three of them were characterized as affinity enhancing in the DMS sets. Gheeraert et al. performed 1 μs long molecular dynamics simulations of five RBD variants (alpha, beta, gamma, delta, and epsilon) in complex with ACE2[13]. They found that L452R, T478K (delta variant) and N501Y (alpha, gamma variants) cause drastic structural changes across the RBD-ACE2 interface. These mutations were classified as binding enriching in the RBD DMS set.

**Table 1.**
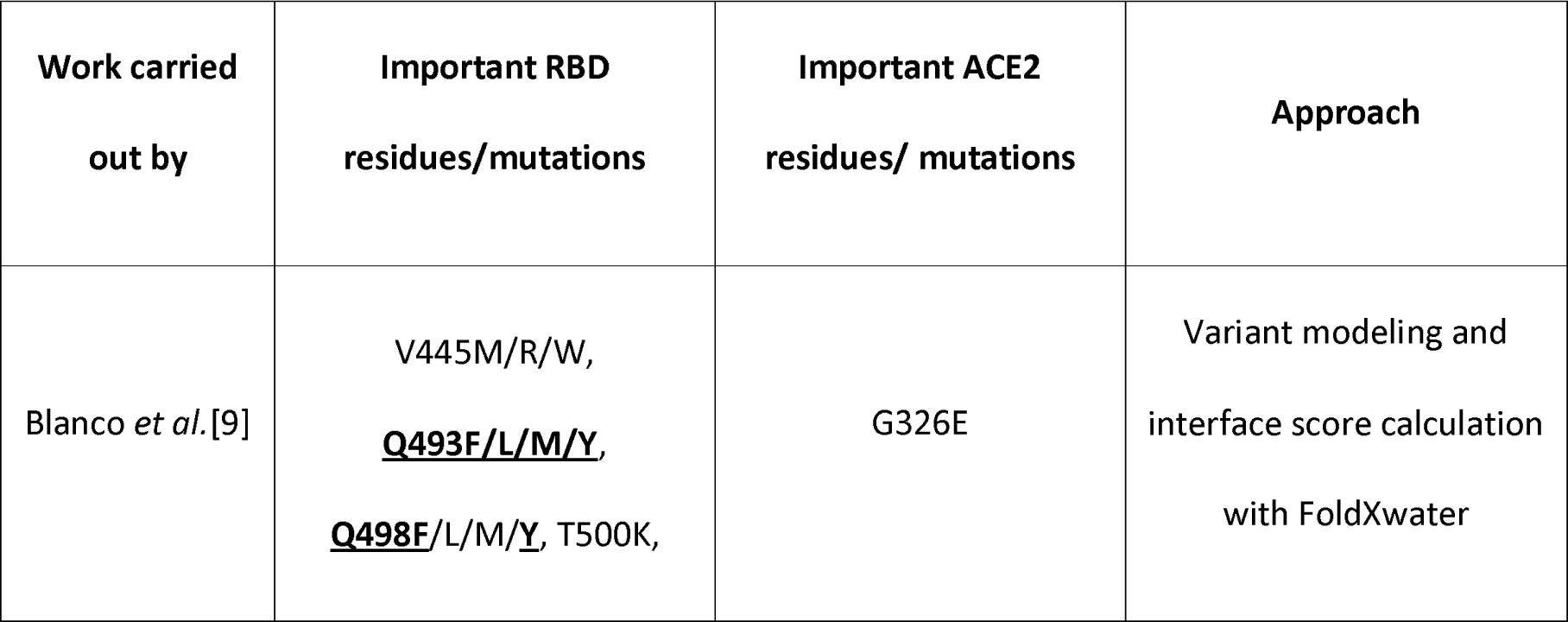
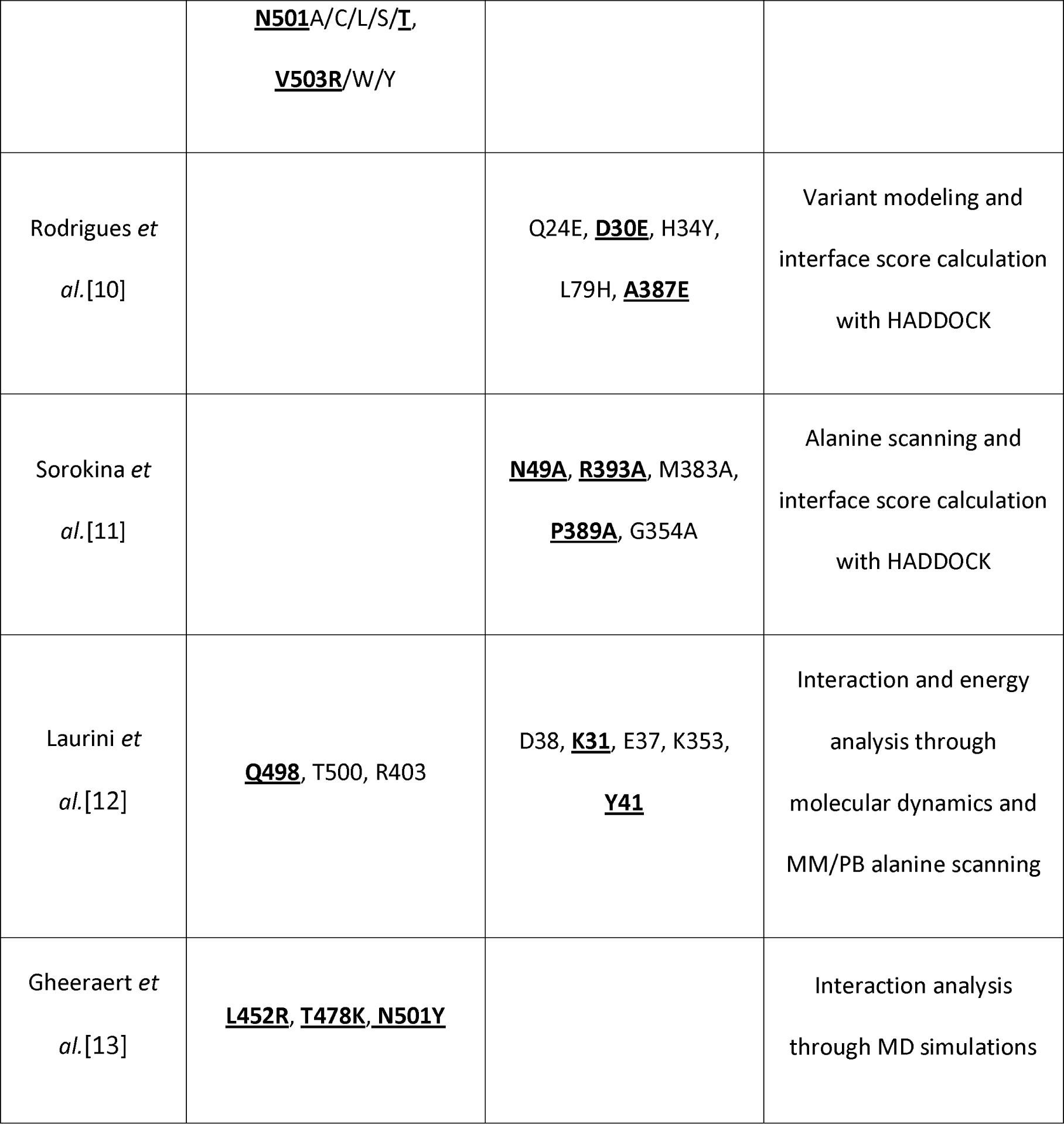
Affinity impacting RBD and ACE2 variant/hotspot predictions. The predictions agreeing with the experimental are underlined and shown in bold.

As presented in Table 1, accurate reproduction of RBD-ACE2 DMS profiles can be tricky even when elaborate simulation techniques are used. So, if such time-intensive simulation approaches are facing challenges in back-calculating the impact of RBD-ACE2 interface variation, how far are the fast prediction tools that were heavily used in the early months of pandemic, such as FoldX and HADDOCK, from accurately predicting the impact of RBD-ACE2 variations? To answer this question, we benchmarked six fast affinity prediction tools, i.e., FoldX, HADDOCK, EvoEF1, MutaBind2, SSIPe, and UEP [14–20] against the Spike- ACE2 interface DMS set. These predictors were selected based on their user-friendliness and speed, since we wanted to put an emphasis on the accessibility of these tools to the researchers who may not have programming experience or enough computing resources. Among these tools, FoldX and EvoEF1 use intra- and inter-molecular energies derived from empirical force field terms. HADDOCK scores complexes by combining intermolecular van der Waals, electrostatics, and empirical desolvation terms. Mutabind and SSPIe utilize FoldX and EvoEF1, respectively, to model the mutations. Both methods have their own scoring functions to consider evolutionary-based information too. UEP scores mutations based on statistically determined intermolecular contact potentials. HADDOCK, MutaBind2, and SSIPe can be run through a web service. FoldX can be called over a GUI through a YASARA plugin. EvoEF1 and UEP are available as stand-alone packages. On top of these conventional tools, we also tested two AI approaches, mmCSM-PPI[21,22] and TopNetTree[23,24] to investigate the impact of AI use in predicting RBD-ACE2 interaction changes. Our benchmarking files can be accessed at https://github.com/CSB-KaracaLab/RBD-ACE2-MutBench with the option to visualize our mutant models at https://rbd-ace2-mutbench.github.io/

## RESULTS and DISCUSSION

### Benchmark Compilation

The deep mutational scanning (DMS) experiments performed by Chan et al. and Starr et al. [7,8] scan the impact of all possible amino acid variations imposed on the RBD domain of Spike and the catalytic domain of ACE2 on RBD-ACE2 binding. These studies classified mutations as binding enriching or depleting when compared to the wild type interactions. While compiling our benchmark set, our aim was (i) to select the DMS subset reporting on the variations across RBD-ACE2 interface, since the selected methods are tuned to predict the impact of interface mutations, (ii) to include an equal number of binding enriching and depleting cases to obtain a balanced benchmark set.

The DMS sets contained 988 interfacial mutations, measured over 26 RBD and 26 ACE2 residues (calculated by PDBePISA[25] on 6m0j PDB[26]). 13% of these 988 mutations were profiled as binding enriching (42 for RBD and 89 for ACE2, Figure 1A) and the rest as binding depleting. As shown in Figure 1A, the binding enriching RBD mutations span a narrow enrichment range [0.01, 0.30], while for ACE2 this range increases to [0.03, 3.37]. We added all enriching cases into our benchmark. To fairly represent the depleting cases, we selected 131 mutations sampling the whole depleting data spread (Figure 1A). We then analyzed the individual binding profiles of selected mutations with heatmaps (Figure 1B). As can be observed from these heatmaps, on the RBD side, several mutations on Q493, S477, F490, N501, V503, E484, Q498 lead to better RBD-ACE2 binding (Figure 1B, Figure S1, Table S1). Among these, Q493R and S477N were observed in omicron; E484K in beta, gamma, eta, iota, mu; E484Q in kappa; N501Y in alpha, beta, gamma, mu, omicron variants[6]. On the ACE2 surface, the top enriching mutations came from T27, Q42, S19, and L79 positions (Figure 1C, Figure S1). All these residues, except S19, were reported as species-associated variations[27]. While appearing less frequently as binding enhancers, K31, E35, M82, and Y83 were earlier listed as critical residues for RBD-ACE2 interactions (Figure 1B-C and Figure S1)[9,12,26]. All these residue positions are situated around the core and the rim of the RBD-ACE2 interface. The top enriching mutations on the RBD side are Q493M, S477D, F490K, N501F, V503M, E484R, Q498H, and on the ACE2 side are T27L, Q42C, S19P (Table S1). Further investigation of these mutations did not lead to a generalized pattern for understanding RBD-ACE2 recognition.

**Figure 1.**
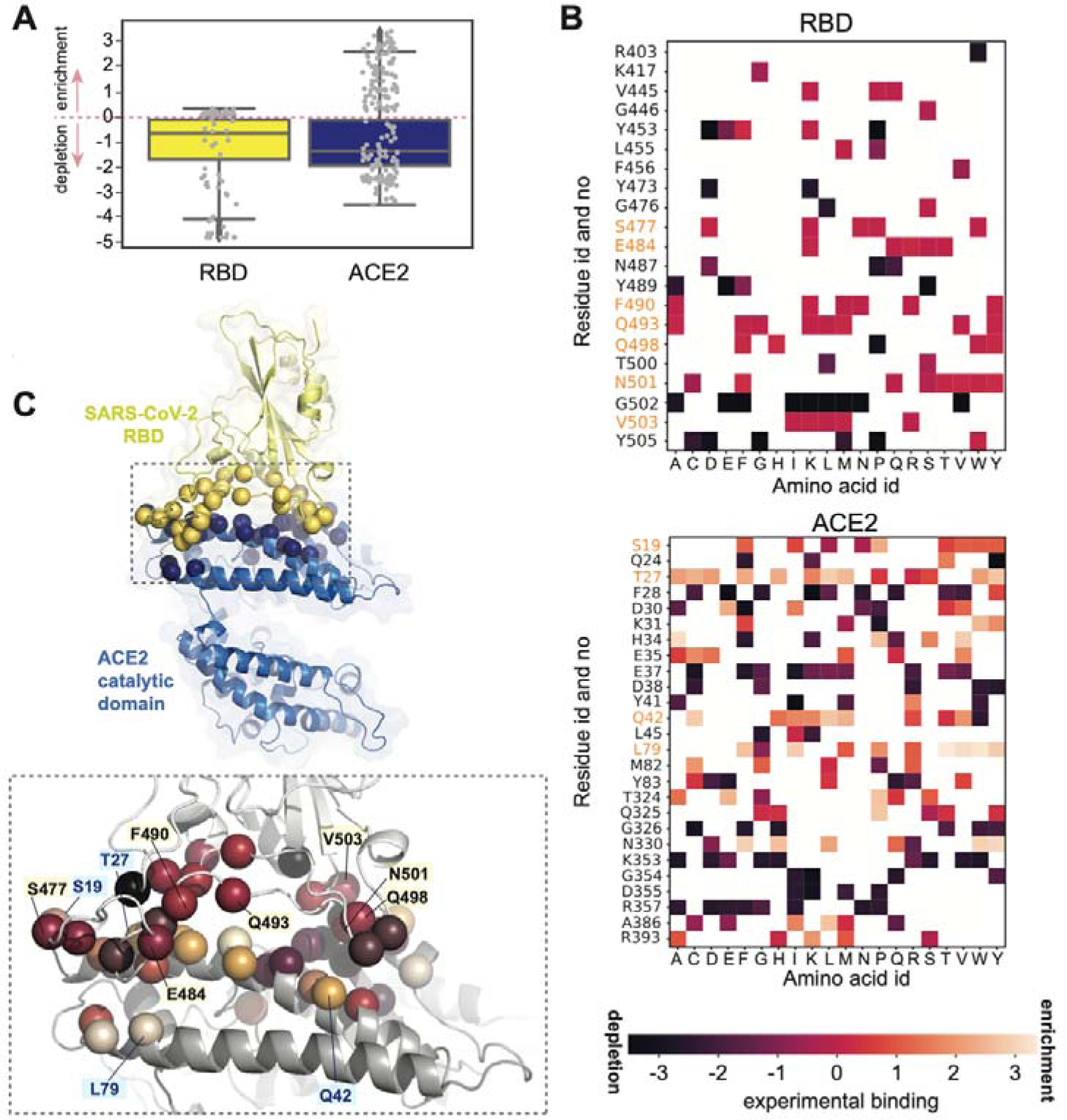
(A) RBD-ACE2 DMS benchmark set. The experimental binding profile distributions of the interfacial RBD (yellow, n=494) and ACE2 (blue, n=494) mutations are represented with box-and-whisker plots. Values >0 indicate binding enriching and values <0 indicate binding depleting mutations. 26.5% of this data set is selected as our benchmark set (n=262): n=84 for RBD (42 enriching, 42 depleting) and n=178 for ACE2 mutations (89 enriching, 89 depleting cases)), represented as gray dots. This panel was generated in Python 3.8 by using pandas, Numpy and Seaborn libraries[34–39]. (B) The experimental binding enrichment and depletion values of our benchmark set, RBD (top) and ACE2 (bottom). The values > 0 correspond to binding enriching positions (light orange), while the values <0 represent the depleting ones (dark purple). The positions leading often to binding enriching mutations are highlighted in orange. (C) The structural depiction of enriching mutations. The interface residues of RBD-ACE2 complex are shown in spheres. The color code of the spheres follows the largest binding value measured for a given residue, as shown in Figure 1B. The important positions are highlighted with labels (yellow: RBD, blue: ACE2). The illustration is generated in PyMOL[28] using PDB 6M0J[26].

### Benchmarking the affinity predictors: seeking for a linear correlation

For all the mutations in our benchmark set (n=262), we calculated the score change imposed by the mutations with FoldX, FoldXwater, EvoEF1, MutaBind2, SSIPe, and HADDOCK. [14–20] (Figure S2). Then, we investigated whether there is any meaningful correlation with the calculated score changes and experimental binding enrichment/depletion values (Figure 2A, Table S2). Here, a perfect correlation would have an absolute value of 1. As a result, we observed insignificant linear correlations for HADDOCK, EvoEF1, and SSIPe predictors. Mediocre correlations with R values ranging from - 0.51 to -0.45 were observed for FoldX and MutaBind2. The best correlation (highest R-value) was obtained by FoldX (R = -0.51). Interestingly, including the water effect into FoldX predictions by using FoldXwater did not improve the accuracy of the original approach (R = - 0.45 vs. R = -0.51). Furthermore, the enhanced scoring function of MutaBind2 built upon FoldX did not improve the original FoldX scoring (R = -0.49 vs. R = -0.51). The same was observed for SSIPe, since it was built upon EvoEF1’s sampling (R = -0.28 vs. -0.27). As the naïve predictor, we ran UEP on a subset of our benchmark (n=129), as UEP is tuned to predict mutation-induced changes of residues that are in contact with more than two atoms. This effort also resulted in an insignificant correlation (R=0.14).

**Figure 2.**
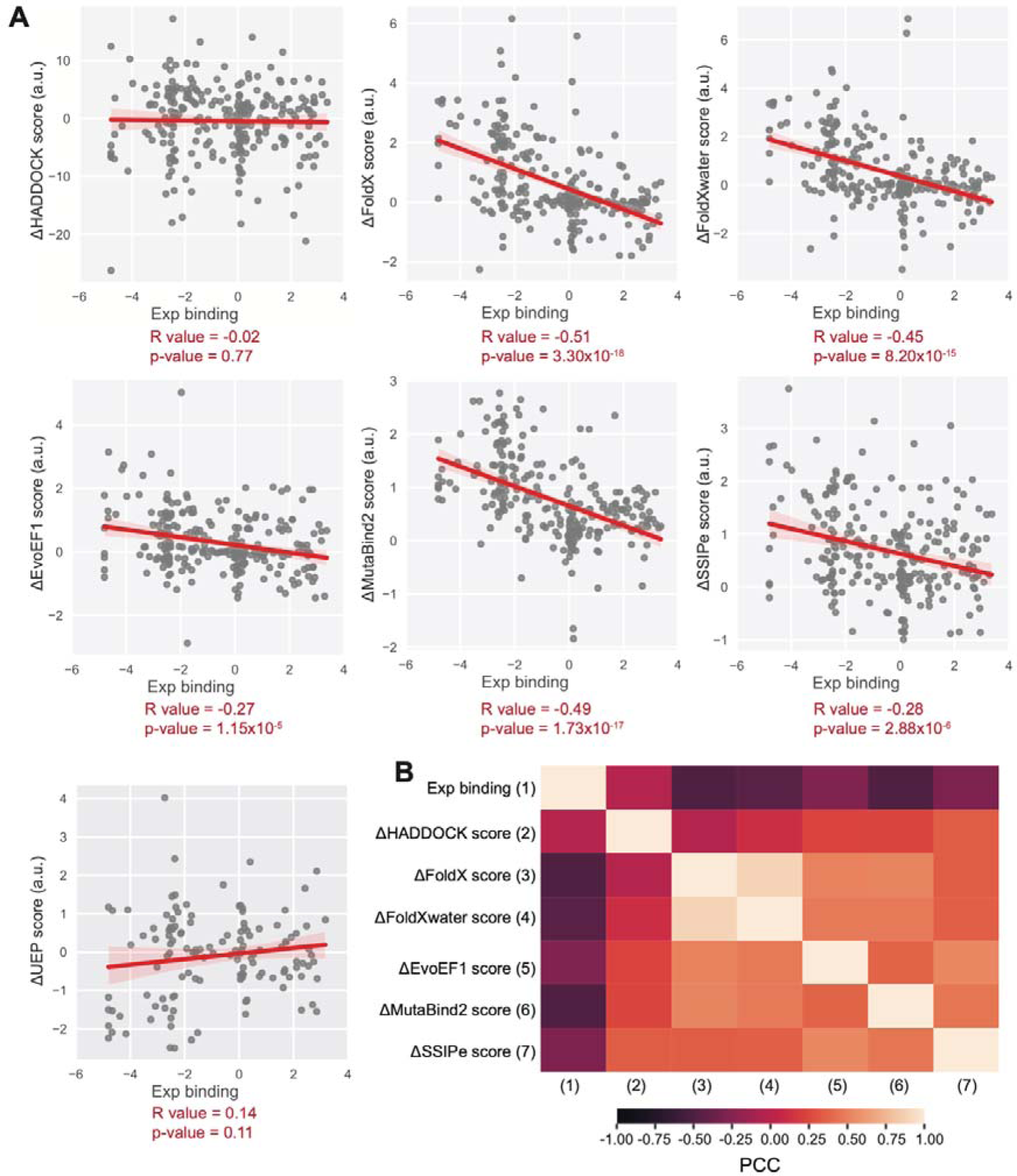
(A) Correlations between experimental DMS benchmark set and the binding affinity predictors. Data points for all scenarios are n=262, except UEP where the number of points is 129 (∼50% of the benchmark set). R- and p-values were calculated by using statistics and scipy libraries of Python 3.8. The statistical data (R and p values) are tabulated in Table S2. (B) The correlation heatmap of the computational and the experimental scores. The correlation values are expressed in terms of Pearson Correlation Coefficients (PCC), ranging between -1 and 1. Highly negatively (PCC = -1) and positively (PCC = 1) correlated predictors are colored with dark purple and light orange, respectively.

We further calculated the pairwise correlations of score changes predicted by each algorithm (Figure 2B). This comparison revealed that HADDOCK reports the most distinct scores compared the other algorithms. We then computed all-atom Root Mean Square Deviations (RMSDs) of each generated mutant model in an all-to-all fashion to understand whether the distinct behavior of HADDOCK came from differentially modeled side chain formations (Figure S3). This analysis demonstrated that HADDOCK indeed generates the largest RMSD models compared to the models computed by the other tools. Notably, MutaBind2 and FoldX conformers resulted in the second highest RMSD cases, indicating that the further minimization steps used by MutaBind2 significantly impacts the final conformation of the FoldX models. As expected, EvoEF1 and SSIPe mutant models came out to be identical, since SSIPe utilizes EvoEF1 to structurally model the mutations.

### Benchmarking the affinity predictors: seeking for a binary classification

Apart from analyzing the correlation between ΔScores and experimental binding values, we also performed a binary assessment. In this assessment, the tools were tested whether they can predict the direction of binding affinity change (as enriching or depleting). Accordingly, we counted a prediction as successful if the experimental and computational data would agree whether a mutation is enriching or depleting. In this regard, the overall prediction accuracy was calculated as the percentage of correct predictions (the success rate). According to this binary assessment, the overall success rates of predictors varied between 54% and 64% (Figure 3A), where the top-ranking predictor once again came out to be FoldX (64%). FoldXwater ranked the second, once again implying that the inclusion of water effects did not improve the prediction accuracy. When we analyzed ACE2 and RBD subsets individually, better prediction rates for depleting mutations were consistently observed, despite the narrow prediction range posed by the experimental data (Figure 3A). Strikingly, MutaBind2 and SSIPe predicted most mutations as depleting, hinting at a problem in using evolutionary-based terms in scoring the host-pathogen system RBD-ACE2. These observations did not change, when we calculated the success rates only for the residues that frequently lead to enriching mutations (the highlighted residues in Figure 1B- C). To investigate this issue further, we calculated the conservation scores of RBD-ACE2 interface amino acids by using ConSurf[29] (Figure S4A). As an outcome, we showed that the ACE2-RBD interface is significantly non-conserved (Figure S4B), which led to the misclassification of most mutation outcomes.

**Figure 3.**
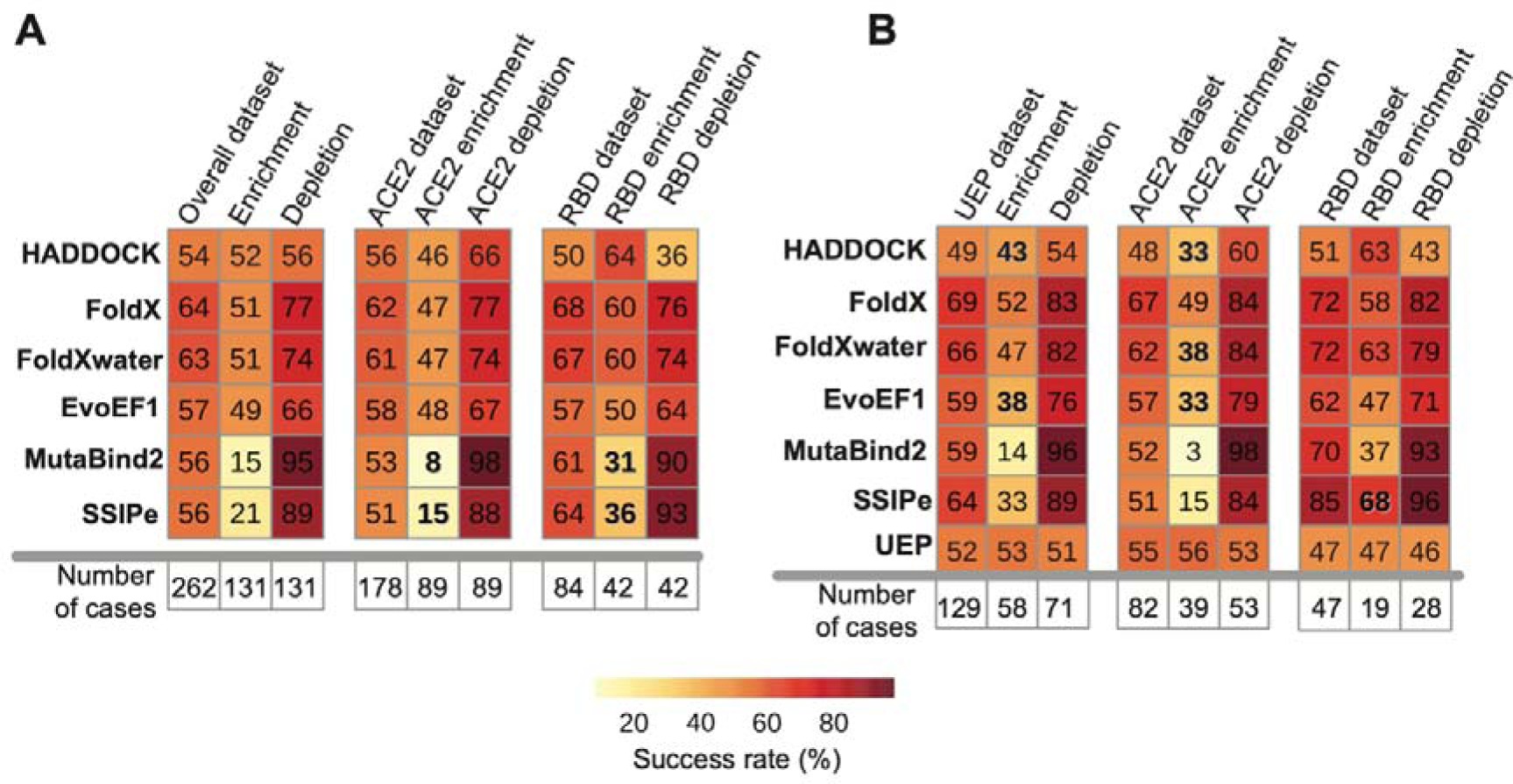
(A) The success rates on the benchmark set (n=262). (B) Success rates of predictors are calculated by using UEP’s data set (n= 129). In both panels, if the success rate changes drastically compared to the overall dataset (first column on the left), it is shown in bold. The plots were prepared by using R-Studio [30–33].

When we further scored the lowest ranking predictor, HADDOCK’s models with FoldX, HADDOCK’s success rate increased by 5% (from 54% from 59%) (Figure S5). Normalizing HADDOCK scores by the buried surface area (BSA) of the interface did not improve the success rates (Figure S5). On the UEP subset, the overall prediction performances vary within a broader range, i.e., 49%-69% (Figure 3B), where the top two predictors became FoldX and FoldXwater (69% vs. 66%) and the lowest performing predictors became UEP and HADDOCK (52% and 49%). On both datasets, all predictors had difficulties in predicting binding-enriching mutations compared to the binding-depleting ones (Figure 3).

### Volume and hydrophobicity biases are the most obvious misprediction determinants

To explore the prediction dependencies of each tool, we assessed their success rates after classifying the benchmark cases according to volume, hydrophobicity, and flexibility changes imposed by the mutations (Figure 4, Table S3, Figure S6). The residue-based volume values were taken from Zhi-hua et al [34], the hydrophobicity ones from Eisenberg et al. [35], and the flexibility ones from Shapovalov and Dunbrack [36]. For each property, we calculated the frequency distributions of the computed changes, given the enrichment or depletion status of the original mutation (Figure 4A-C). To serve as a background, we calculated the frequency distributions of the experimental data too (gray lines in Figure 4A- C). Here our assumption was a discrepancy between the tails of the experimental data and the predictors’ distributions should point to an obvious bias for a given property. Accordingly, when we investigated the volume change plots, we saw that all predictors had difficulties in predicting volume decreasing mutations when the mutation induces an enrichment in binding (Figure 4A). In the case of depleting mutations, only HADDOCK has an apparent bias toward classifying volume increasing mutations wrongly. In the case of hydrophobicity change, for all predictors, there is a slight tendency to classify hydrophobicity decreasing binding enriching mutations wrongly. This is more pronounced for hydrophobicity decreasing and binding depleting mutations for HADDOCK. We did not observe any bias for the predictors regarding the changes in the side chain flexibilities.

**Figure 4.**
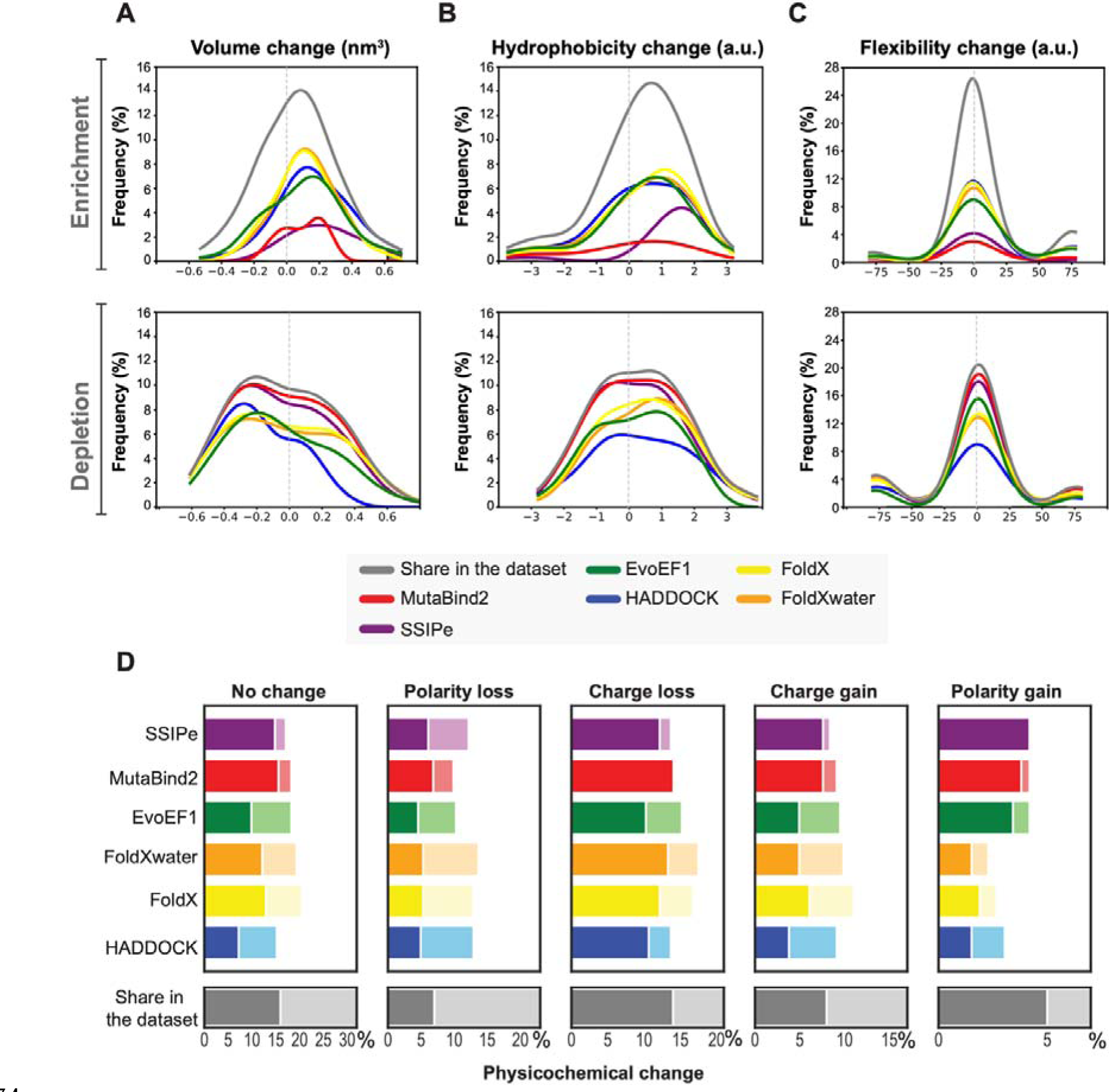
The effects of mutation-induced changes in the amino acid physical properties on the predictor’s success rates according to (A) Volume change, (B) Hydrophobicity change, (C) Flexibility change. The original data set distribution for a given binding class is plotted in gray. The other predictors are colored as provided in the legend. Volume/hydrophobicity/flexibility-increasing mutations reside on the “positive side” of the plot (x-axis value >0), whereas volume/hydrophobicity/flexibility-decreasing mutations reside on the “negative side” of the pot (x- axis value <0) (D) The percentage of the correctly predicted cases, given the physicochemical change induced upon mutation when polarity and charge states are considered. and light colors represent successfully predicted depleting and enriching cases, respectively. The original data share for a given class in the dataset is plotted in grey.

To gauge whether the enriching and depletion mutation predictions were balanced for a given property, we calculated the difference between the area under the curve of the successfully predicted enriching and depletion mutations (ΔSuccess, Figure S6). Here, our assumption was, if the prediction is balanced, then the ΔSuccess should be close to zero. On the other hand, the ΔSuccess values towards 100 and -100 should indicate extreme biases. This analysis revealed that HADDOCK has moderate volume change bias with 34 ΔSuccess score. Normalizing HADDOCK scores by the BSA of the interface doubled this bias leading to 78 ΔSuccess score. When HADDOCK models were scored with FoldX, the original 34 ΔSuccess shifted to -31 ΔSuccess, resulting in a bias toward depleting mutations (Figure S6). When we considered the success rates based on hydrophobicity, we found out that FoldX tends to predict depleting cases better with -31 ΔSuccess score. Inclusion of water in FoldX (FoldXwater) increases this moderate bias (ΔSuccess score -36 vs -31). The bias towards increased hydrophobicity might stem from the fact that only three of the 26 ACE2 interface residues (Y41, Y83, K353) and only six of the 26 RBD interface residues (L455, F456, N487, Y89, Q498, N501) are core interface residues, while the rest are partially or totally solvent exposed (as calculated by EPPIC [37]). The predictors are not tuned to perform well on such unusual binding sites, where a short fraction of the interface is composed of buried hydrophobic residues. As an outcome, we observed challenges in predicting mutations with changes in hydrophobicity. Finally, we could not find any relation between the flexibility change and success rate of predictors (Figure 4C and Figure S6). Finally, we found that ΔSuccess is extremely skewed for all metrics for MutaBind2 and SSIPe, since they predict almost all mutations as depleting.

To present a complete analysis, we also investigated the impact of the physicochemical property changes induced by each mutation on the prediction accuracy (Figure 4D). The physicochemical change classes we considered were no change, polarity loss/gain, and charge loss/gain. To serve as a background, we calculated the share of each class within the original data set (gray bars in Figure 4D). As a result, we could not observe any specific bias given a physicochemical property change. The success shares reflect the general trends observed in successfully predicting binding enriching or depletion mutations, as shown in Figure 4A.

Which other physicochemical factors could play a role in the misprediction of mutations? Interestingly, we could not observe any bias over the correctly predicted cases when the experimental data spared was considered (Figure S7A). This observation directed our attention towards a factor that is missing in all these assessments, namely, the RBD/ACE2 glycan chains. Both the catalytic domain of ACE2 and RBD contains multiple glycosylation sites, the impact of which were explored by detailed all-atom MD simulations [38–44]. Nguyen et al., for example, conducted simulations involving non-glycan, MAN9-glycan, and FA2-glycan ACE2-RBD MD simulations for more than 200 microseconds [38]. As an outcome, they revealed that ACE2 glycans impact virus binding affinity through electrostatic effects, without disrupting the physical contacts established between the virus and its host. The direct impact of the glycans on the ACE2-RBD binding affinity were demonstrated experimentally too[45]. To explore the potential impact of missing glycan chains in our calculations, we computed the distances from wrongly predicted mutation sites to the six ACE2 glycosylation sites (N53, N90, N103, N322, N432, N546) (Figure S7B). This analysis showed that for all the predictors, at least one fourth of the wrongly predicted mutation sites fall within 20 Å of these six sites, endorsing the role of missing glycans. Here, we should note that HADDOCK provides an option to incorporate glycans into the predictions. Though, as demonstrated earlier, its use will be limited to very short glycan chains presented in the reference EM structure[10]. We therefore did not resort to this option for the sake of being consistent in our benchmarking.

### Can AI methods perform better than the classical techniques?

During recent years, several AI methods have been proposed for predicting mutation-imposed interaction changes [21–23,46–48]. Among these tools, we concentrated on two, mmCSM-PPI[22] and TopNetTree[23,24], of which results on the RBD-ACE2 system were readily available. [24,49–56]. The machine learning approach, mmCSM-PPI, utilizes physicochemical and geometrical properties of protein structures within a graph-based structural framework to model the impact of mutations on the inter-residue interaction network. mmCSM-PPI includes evolutionary scores, non-covalent interactions, and dynamics terms from Normal Mode Analysis. We ran mmCSM-PPI against our experimental data set through their user-friendly web interface and obtained an R value of 0.53, which is as high as the one from FoldX (Figure S8A, Table S2). mmCSM-PPI2 produced the most similar results to MutaBind2 and SSIPe, with PCC values of 0.70 and 0.50, respectively (Figure S8B). In predicting the direction of mutation impact, the overall success rate for mmCSM-PPI2 became 57%, with success rates of 22% for enriching and 92% for depleting mutations (Figure S8C-D). The failure of mmCSM-PPI in predicting the enriching mutations aligns well with the behavior of other two evolutionary-inclusive MutaBind2 and SSIPe algorithms.

TopNetTree, a recent deep learning approach, includes physical pairwise interactions, Euclidean distances, and cavity structures within a topological framework. Notably, TopNetTree has been actively used in numerous SARS-CoV-2 studies[46,50,52–54,56]. In particular, Chen et al., trained TopNetTree on SARS-CoV-2 datasets to accurately predict changes in binding free energy for the S protein, ACE2, or antibodies induced by mutations[24]This tool was not available as a web server or a standalone tool. Though, since its RBD mutation profiles were published earlier, we could take them as a basis in our assessment [24]. Over our RBD data set, TopNetTree obtained an R value of -0.01 (Figure S9A), indicating a lack of meaningful correlation. So, although this approach was specifically trained to predict the outcome of mutations on the complete S protein, it fails to predict the impact of its interfacial mutations. We further observed that, like in the other methods, TopNetTree predicts the depleting cases more efficiently (64% success rate) than the enriching ones (48% success rate) (Figure S9C-D). It also generates the most diverse set of predictions compared to the other probed methods (Figure S9B).

Expanding on these results, we claim that, contrary to expectation, machine/deep learning approaches do not yield significantly better results on the RBD-ACE2 system, compared to the classical force-field-based techniques.

## CONCLUSION

In the early months of SARS-CoV-2 pandemic, several fast and user-friendly mutation modeling and scoring tools, such as FoldX and HADDOCK, were heavily used to predict the impact of Spike/ACE2 variations across the Spike-ACE2 interface. Expanding on these efforts, in this work, we benchmarked six fast and commonly used structure-based binding affinity predictors (FoldX, EvoEF1, MutaBind2, SSIPe, HADDOCK, and UEP) and two AI approaches (mCSM-PPI and TopNetTree) against the RBD-ACE2 DMS binding data. As a result, we observed that none of the predictors could produce a meaningful correlation with the experimental data (best correlation R-value -0.51 was obtained with FoldX). Even when a binary classification (binding enriching/depleting) was considered, the highest accuracy was obtained by FoldX with 64% success rate. Furthermore, all predictors had difficulties in predicting binding enriching mutations, especially the ones using conservation-based terms in their scoring. Finally, the most obvious biases in mispredictions were found to be toward volume and hydrophobicity changes, especially for HADDOCK and FoldX, respectively.

These observations suggest plenty of room to improve the affinity predictors for guessing the variant-induced binding profile changes of host-pathogen systems, such as Spike-ACE2. To aid such improvements we provide our benchmarking data at https://github.com/CSB-KaracaLab/RBD-ACE2-MutBench with the option to visualize our mutant models at https://rbd-ace2-mutbench.github.io/. We hope that our benchmarking study will guide the computational community for being prepared not only for combatting SARS-CoV-2-related health concerns but also other infectious diseases.

## MATERIALS and METHODS

### Interfacial DMS value selection

The original DMS sets contained 3,819 and 2,223 point mutations for RBD and ACE2, respectively. From this set, we isolated the interfacial 988 RBD-ACE2 point mutations, for the following residues (calculated over 6m0j[26] with PDBePISA[25] (Figure 1C)):

*Twenty-six RBD positions: R403, K417, V445, G446, Y449, Y453, L455, F456, Y473, A475, G476, S477, Q484, G485, F486, N487, Y489, F490, Q493, G496, Q498, T500, N501, G502, V503, and Y505*.

*Twenty-six ACE2 positions: S19, Q24, T27, F28, D30, K31, H34, E35, E37, D38, Y41, Q42, L45, L79, M82, Y83, T324, Q325, G326, N330, K353, G354, D355, R357, A386, R393*.

### Structure-based binding affinity predictors

Below are the brief scoring description of each predictor used. Except for UEP, all predictors explicitly model the mutation according to predictor’s force field. Except for HADDOCK, these predictors sample a single conformation of the mutation.

FoldX: The scoring function of FoldX is a linear sum of vdW energy (ΔG_vdw_), hydrophobic (ΔG_solvH_) and polar group desolvation (ΔG_solvP_) energies, hydrogen bond energy from water molecules (ΔG_wb_), hydrogen bond energy (ΔG_hbond_), electrostatic energy (ΔG_el_), the electrostatic contribution of different polypeptides (ΔGk_on_), entropic penalty for backbone (ΔS_mc_), entropic penalty of the side chain (ΔS_sc_), and steric overlaps (ΔG_clash_) (Eq. 1). FoldX also has an option to include the contribution of water molecules to the binding affinity (FoldXwater).

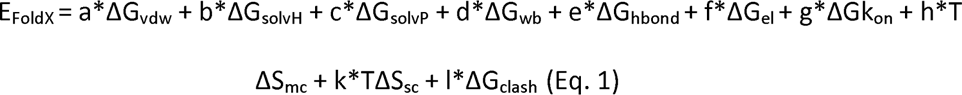

EvoEF1: The scoring function of EvoEF1 contains van der Waals (E_vdw_), electrostatics (E_elec_), hydrogen bond (E_HB_), desolvation energies (E_solv_), and the energy of a reference state (E_ref_) (Eq. 2).

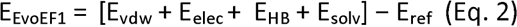

MutaBind2 and SSIPe: use FoldX and EvoEF1, respectively, to explicitly model the desired mutation. MutaBind2 further imposes relaxation and utilizes extra force field and contact-based terms, together with a metric measuring the evolutionary conservation of the mutation site. All these terms are incorporated into a random forest based scoring algorithm. SSIPe uses EvoEF1 energy terms and residue conservation-related terms, extracted from iAlign[57] and PSI-BLAST[58].

HADDOCK: In this work, we used HADDOCK water refinement to model the mutations. The complexes then scored according to the sum of three terms, van der Waals, electrostatic, and desolvation energy (Eq.3) [59]. For each HADDOCK modeling, we generated 250 conformations, and subsequently selected the conformation with the lowest HADDOCK score for further analysis.

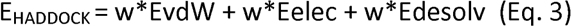

UEP: UEP predicts the impact of all possible interfacial mutations, when the position of interest has interactions with at least two other residues (within 5Å range). The scoring function of UEP expands on the statistically determined intermolecular contact potentials.

To run FoldX and EvoEF1, we used their stand-alone packages (http://foldxsuite.crg.eu/products#foldx, https://github.com/tommyhuangthu/EvoEF). HADDOCK, MutaBind2, and SSIPe were run on their servers, as given in

https://milou.science.uu.nl/services/HADDOCK2.2/, https://lilab.jysw.suda.edu.cn/research/mutabind2/ https://zhanggroup.org/SSIPe/. MutaBind2, SSIPe, and UEP directly provide the binding affinity change predictions, whereas for the rest we calculated the predicted binding affinity change according to Eq. 4:

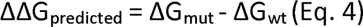

A mutation is evaluated as binding enriching if the predicted binding value change (ΔScore_predicted_) is <0 and binding depleting, if (ΔScore_predicted_) is >0.

### Conservation analysis

To investigate why the evolutionary-based approaches failed, we used ConSurf[29] on RBD and ACE2. ConSurf assigns a conservation score to each residue within the protein complex, ranging from 1 to 9, with 1 indicating non-conserved residues and 9 signifying highly conserved ones.

### Performance evaluation according to change in amino acid physical properties upon a mutation

The predictions were evaluated from the perspectives of volume, hydrophobicity, flexibility, and physicochemical property change upon mutation (ΔProperty_change_ = Property_mutation_ - Property_wildtype_, Table S3). The physicochemical properties considered were: polar amino acids - N, Q, S, T, Y; non-polar amino acids - A, G, I, L, M, F, P, W, V, C; charged amino acids - H, E, D, R, K. Success rate and metric evaluations were performed in Python 3.8.5 with Pandas, Numpy, seaborn, and Matplotlib libraries [60–65]. For each category, the percentage of successfully predicted cases were calculated by Eq .5.

Success rate = Correct-Predictions/All-Predictions*100 (Eq. 5)

## DATA AVAILABILITY

All results including the codes and notebooks are deposited in Github (https://github.com/CSB-KaracaLab/RBD-ACE2-MutBench) and the models and the scores can be visualized at https://rbd-ace2-mutbench.github.io/.

## SUPPLEMENTARY INFORMATION

Supplementary Information are submitted with the manuscript.

## ACKNOWLEDGEMENTS

All the simulations and analyses were carried out in the HPC resources of Izmir Biomedicine and Genome Center. The authors would like to thank João P. G. L. M. Rodrigues for the critical reading of our manuscript and providing help in building our results visualization page.

## AUTHOR CONTRIBUTIONS

B.O. performed predictor runs, statistical analysis and RMSD calculations, generated structural and statistical figures, edited and finalized all the figures, prepared the Github page, and wrote the manuscript. E. Ş. performed predictor runs, calculated the physical property and binary success rates, prepared the Github page, contributed to writing Introduction and Methods & Materials sections. A.Ö performed deep learning predictor runs and the statistical analysis. M.E. generated heatmap tables for success rate results and wrote the manuscript. M.K. performed predictor runs, generated heatmaps for the RMSD calculations and predictor scores. M.O. performed the most enriching mutation analysis. C.Y. performed predictor runs. N. A. and G. K. prepared the visualization page on Github. A. B. B. and B.S. worked on the conceptualization of the benchmark set. E. K. conceptualized the study, overlooked the project, and wrote the manuscript.

## REFERENCES

[1] LeDuc JW, Barry MA. SARS, the First Pandemic of the 21st Century. Emerg Infect Dis 2004;10:e26. 10.3201/EID1011.040797_02.

[2] Durai P, Batool M, Shah M, Choi S. Middle East respiratory syndrome coronavirus: transmission, virology and therapeutic targeting to aid in outbreak control. Exp Mol Med 2015;47:e181. 10.1038/EMM.2015.76.

[3] Platto S, Xue T, Carafoli E. COVID19: an announced pandemic n.d. 10.1038/s41419-020-02995-9.

[4] Li W, Moore MJ, Vasllieva N, Sui J, Wong SK, Berne MA, et al. Angiotensin- converting enzyme 2 is a functional receptor for the SARS coronavirus. Nature 2003;426:450. 10.1038/NATURE02145.

[5] Ali A, Vijayan R. Dynamics of the ACE2–SARS-CoV-2/SARS-CoV spike protein interface reveal unique mechanisms. Sci Rep 2020;10. 10.1038/s41598-020-71188-3.

6. CoVariants n.d. https://covariants.org/ (accessed June 27, 2022).

[7] Chan KK, Dorosky D, Sharma P, Abbasi SA, Dye JM, Kranz DM, et al. Engineering human ACE2 to optimize binding to the spike protein of SARS coronavirus 2. Science (1979) 2020;369. 10.1126/SCIENCE.ABC0870.

[8] Starr TN, Greaney AJ, Hilton SK, Ellis D, Crawford KHD, Dingens AS, et al. Deep Mutational Scanning of SARS-CoV-2 Receptor Binding Domain Reveals Constraints on Folding and ACE2 Binding. Cell 2020;182:1295–1310.e20. 10.1016/j.cell.2020.08.012.

[9] Delgado Blanco J, Hernandez-Alias X, Cianferoni D, Serrano L. In silico mutagenesis of human ACE2 with S protein and translational efficiency explain SARS-CoV-2 infectivity in different species. PLoS Comput Biol 2020;16:e1008450. 10.1371/journal.pcbi.1008450.

[10] Rodrigues JPGLM, Barrera-Vilarmau S, M. C. Teixeira J, Sorokina M, Seckel E, Kastritis PL, et al. Insights on cross-species transmission of SARS-CoV-2 from structural modeling. PLoS Comput Biol 2020;16:e1008449. 10.1371/journal.pcbi.1008449.

[11] Sorokina M, M. C. Teixeira J, Barrera-Vilarmau S, Paschke R, Papasotiriou I, Rodrigues JPGLM, et al. Structural models of human ACE2 variants with SARS-CoV-2 Spike protein for structure-based drug design. Sci Data 2020;7:309. 10.1038/s41597-020-00652-6.

[12] Laurini E, Marson D, Aulic S, Fermeglia M, Pricl S. Computational Alanine Scanning and Structural Analysis of the SARS-CoV-2 Spike Protein/Angiotensin-Converting Enzyme 2 Complex. ACS Nano 2020;14. 10.1021/acsnano.0c04674.

[13] Gheeraert A, Vuillon L, Chaloin L, Moncorgé O, Very T, Perez S, et al. Singular Interface Dynamics of the SARS-CoV-2 Delta Variant Explained with Contact Perturbation Analysis. J Chem Inf Model 2022;62:3107–22. 10.1021/ACS.JCIM.2C00350.

[14] Schymkowitz J, Borg J, Stricher F, Nys R, Rousseau F, Serrano L. The FoldX web server: An online force field. Nucleic Acids Res 2005;33. 10.1093/nar/gki387.

[15] Pearce R, Huang X, Setiawan D, Zhang Y. EvoDesign: Designing Protein– Protein Binding Interactions Using Evolutionary Interface Profiles in Conjunction with an Optimized Physical Energy Function. J Mol Biol 2019;431. 10.1016/j.jmb.2019.02.028.

[16] Zhang N, Chen Y, Lu H, Zhao F, Alvarez RV, Goncearenco A, et al. MutaBind2: Predicting the Impacts of Single and Multiple Mutations on Protein-Protein Interactions. IScience 2020;23. 10.1016/j.isci.2020.100939.

[17] Huang X, Zheng W, Pearce R, Zhang Y, Zhang Y. SSIPe: Accurately estimating protein-protein binding affinity change upon mutations using evolutionary profiles in combination with an optimized physical energy function. Bioinformatics 2020;36. 10.1093/bioinformatics/btz926.

[18] Van Zundert GCP, Rodrigues JPGLM, Trellet M, Schmitz C, Kastritis PL, Karaca E, et al. The HADDOCK2.2 Web Server: User-Friendly Integrative Modeling of Biomolecular Complexes. J Mol Biol 2016. 10.1016/j.jmb.2015.09.014.

[19] Honorato R V., Koukos PI, Jiménez-García B, Tsaregorodtsev A, Verlato M, Giachetti A, et al. Structural Biology in the Clouds: The WeNMR-EOSC Ecosystem. Front Mol Biosci 2021;8. 10.3389/fmolb.2021.729513.

[20] Amengual-Rigo P, Fernández-Recio J, Guallar V. UEP: an open-source and fast classifier for predicting the impact of mutations in protein-protein complexes. Bioinformatics 2021;37. 10.1093/bioinformatics/btaa708.

[21] Rodrigues CHM, Pires DEV, Ascher DB. MmCSM-PPI: Predicting the effects of multiple point mutations on protein-protein interactions. Nucleic Acids Res 2021;49:W417–24. 10.1093/nar/gkab273.

[22] Rodrigues CHM, Myung Y, Pires DEV, Ascher DB. MCSM-PPI2: predicting the effects of mutations on protein-protein interactions. Nucleic Acids Res 2019;47:W338–44. 10.1093/nar/gkz383.

[23] Wang M, Cang Z, Wei GW. A topology-based network tree for the prediction of protein–protein binding affinity changes following mutation. Nat Mach Intell 2020;2:116–23. 10.1038/s42256-020-0149-6.

[24] Chen J, Wang R, Wang M, Wei GW. Mutations Strengthened SARS-CoV-2 Infectivity. J Mol Biol 2020;432:5212–26. 10.1016/j.jmb.2020.07.009.

[25] Krissinel E, Henrick K. Inference of Macromolecular Assemblies from Crystalline State. J Mol Biol 2007;372:774–97. 10.1016/j.jmb.2007.05.022.

[26] Lan J, Ge J, Yu J, Shan S, Zhou H, Fan S, et al. Structure of the SARS-CoV-2 spike receptor-binding domain bound to the ACE2 receptor. Nature 2020;581:215–20. 10.1038/s41586-020-2180-5.

[27] Wan Y, Shang J, Graham R, Baric RS, Li F. Receptor Recognition by the Novel Coronavirus from Wuhan: an Analysis Based on Decade-Long Structural Studies of SARS Coronavirus. J Virol 2020;94. 10.1128/jvi.00127-20.

[28[ Schrödinger LLC and DW. PyMOL n.d.

[29] Ashkenazy H, Abadi S, Martz E, Chay O, Mayrose I, Pupko T, et al. ConSurf 2016: an improved methodology to estimate and visualize evolutionary conservation in macromolecules. Nucleic Acids Res 2016;44:W344–50. 10.1093/NAR/GKW408.

[30[ Integrated Development Environment for R. RStudio n.d.

[31[ A Language and Environment for Statistical Computing. R n.d.

[32] Kolde R. Pheatmap: Pretty Heatmaps 2019.

[33] Neuwirth E. RColorBrewer: ColorBrewer Palettes 2014.

[34] Lin Z, Long H, Bo Z, Wang Y, Wu Y. New descriptors of amino acids and their application to peptide QSAR study. Peptides (NY) 2008;29:1798–805. 10.1016/j.peptides.2008.06.004.

[35] Eisenberg D, Schwarz E, Komaromy M, Wall R. Analysis of membrane and surface protein sequences with the hydrophobic moment plot. J Mol Biol 1984;179:125–42. 10.1016/0022-2836(84)90309-7.

[36] Shapovalov MV, Dunbrack RL. A Smoothed Backbone-Dependent Rotamer Library for Proteins Derived from Adaptive Kernel Density Estimates and Regressions. Structure 2011;19:844–58. 10.1016/j.str.2011.03.019.

[37] Bliven S, Lafita A, Parker A, Capitani G, Duarte JM. Automated evaluation of quaternary structures from protein crystals. PLoS Comput Biol 2018;14. 10.1371/journal.pcbi.1006104.

[38] Nguyen K, Chakraborty S, Mansbach RA, Korber B, Gnanakaran S. Exploring the role of glycans in the interaction of sars-cov-2 rbd and human receptor ace2. Viruses 2021;13. 10.3390/v13050927.

[39] Mehdipour AR, Hummer G. Dual nature of human ACE2 glycosylation in binding to SARS-CoV-2 spike 2021;118:2100425118. 10.1073/pnas.2100425118/-/DCSupplemental.

[40] Zhao P, Praissman JL, Grant OC, Cai Y, Xiao T, Rosenbalm KE, et al. Virus- Receptor Interactions of Glycosylated SARS-CoV-2 Spike and Human ACE2 Receptor. Cell Host Microbe 2020;28:586–601.e6. 10.1016/j.chom.2020.08.004.

[41] Gong Y, Qin S, Dai L, Tian Z. The glycosylation in SARS-CoV-2 and its receptor ACE2. Signal Transduct Target Ther 2021;6. 10.1038/s41392-021-00809-8.

[42] Mugnai ML, Shin S, Thirumalai D. Entropic contribution of ACE2 glycans to RBD binding. Biophys J 2023;122:2506–17. 10.1016/j.bpj.2023.05.003.

[43] Cao W, Dong C, Kim S, Hou D, Tai W, Du L, et al. Biomechanical characterization of SARS-CoV-2 spike RBD and human ACE2 protein-protein interaction. Biophys J 2021;120:1011–9. 10.1016/j.bpj.2021.02.007.

[44] Acharya A, Lynch DL, Pavlova A, Pang YT, Gumbart JC. ACE2 glycans preferentially interact with SARS-CoV-2 over SARS-CoV. Chemical Communications 2021;57:5949–52. 10.1039/D1CC02305E.

[45] Isobe A, Arai Y, Kuroda D, Okumura N, Ono T, Ushiba S, et al. ACE2 N- glycosylation modulates interactions with SARS-CoV-2 spike protein in a site- specific manner. Commun Biol 2022;5. 10.1038/s42003-022-04170-6.

[46] Mohseni Behbahani Y, Laine E, Carbone A. Deep Local Analysis deconstructs protein-protein interfaces and accurately estimates binding affinity changes upon mutation. Bioinformatics 2023;39:I544–52. 10.1093/bioinformatics/btad231.

[47] Yue Y, Li S, Wang L, Liu H, Tong HHY, He S. MpbPPI: a multi-task pre- training-based equivariant approach for the prediction of the effect of amino acid mutations on protein–protein interactions. Brief Bioinform 2023. 10.1093/bib/bbad310.

[48] Rodrigues CHM, Pires DEV, Ascher DB. DynaMut2: Assessing changes in stability and flexibility upon single and multiple point missense mutations. Protein Science 2021;30:60–9. 10.1002/pro.3942.

[49] Xiong Q, Cao L, Ma C, Tortorici MA, Liu C, Si J, et al. Close relatives of MERS-CoV in bats use ACE2 as their functional receptors. Nature 2022;612:748–57. 10.1038/s41586-022-05513-3.

[50] Chen J, Gao K, Wang R, Wei GW. Revealing the Threat of Emerging SARS-CoV-2 Mutations to Antibody Therapies. J Mol Biol 2021;433. 10.1016/j.jmb.2021.167155.

[51] Khan A, Gui J, Ahmad W, Haq I, Shahid M, Khan AA, et al. The SARS-CoV-2 B.1.618 variant slightly alters the spike RBD-ACE2 binding affinity and is an antibody escaping variant: a computational structural perspective. RSC Adv 2021;11:30132–47. 10.1039/d1ra04694b.

[52] Chen J, Wang R, Gilby NB, Wei G-W. Omicron Variant (B.1.1.529): Infectivity, Vaccine Breakthrough, and Antibody Resistance. J Chem Inf Model 2022;62:412–22. 10.1021/acs.jcim.1c01451.

[53] Shan S, Luo S, Yang Z, Hong J, Su Y, Ding F, et al. Deep learning guided optimization of human antibody against SARS-CoV-2 variants with broad neutralization. Proceedings of the National Academy of Sciences 2022;119. 10.1073/pnas.2122954119.

[54] Wang R, Chen J, Gao K, Hozumi Y, Yin C, Wei GW. Analysis of SARS-CoV- 2 mutations in the United States suggests presence of four substrains and novel variants. Commun Biol 2021;4. 10.1038/s42003-021-01754-6.

[55] Wang R, Chen J, Gao K, Wei G-W. Vaccine-escape and fast-growing mutations in the United Kingdom, the United States, Singapore, Spain, India, and other COVID-19-devastated countries. Genomics 2021;113:2158–70. 10.1016/j.ygeno.2021.05.006.

[56] Zeng L, Lu Y, Yan W, Yang Y. A Protein Co-Conservation Network Model Characterizes Mutation Effects on SARS-CoV-2 Spike Protein. Int J Mol Sci 2023;24:3255. 10.3390/ijms24043255.

[57] Gao M, Skolnick J. iAlign: a method for the structural comparison of protein– protein interfaces. Bioinformatics 2010;26:2259–65. 10.1093/bioinformatics/btq404.

[58] Altschul SF, Madden TL, Schäffer AA, Zhang J, Zhang Z, Miller W, et al. Gapped BLAST and PSI-BLAST: A new generation of protein database search programs. Nucleic Acids Res 1997;25. 10.1093/nar/25.17.3389.

[59] Dominguez C, Boelens R, Bonvin AMJJ. HADDOCK: A Protein−Protein Docking Approach Based on Biochemical or Biophysical Information. J Am Chem Soc 2003;125:1731–7. 10.1021/ja026939x.

[60] Van Rossum G, Drake FL, Harris CR, Millman KJ, van der Walt SJ, Gommers R, et al. Python 3 Reference Manual. vol. 585. 2009.

[61] Harris CR, Millman KJ, van der Walt SJ, Gommers R, Virtanen P, Cournapeau D, et al. Array programming with NumPy. Nature 2020;585. 10.1038/s41586-020-2649-2.

[62] Kluyver T, Ragan-Kelley B, Pérez F, Granger B, Bussonnier M, Frederic J, et al. Jupyter Notebooks—a publishing format for reproducible computational workflows. Positioning and Power in Academic Publishing: Players, Agents and Agendas - Proceedings of the 20th International Conference on Electronic Publishing, ELPUB 2016, 2016. 10.3233/978-1-61499-649-1-87.

[63] Anaconda Software Distribution. Anaconda Documentation 2020.

[64] The pandas development team. Pandas-dev/pandas: Pandas. Zenodo 2020.

[65] Waskom M. seaborn: statistical data visualization. J Open Source Softw 2021;6. 10.21105/joss.03021.

