## Supporting Information for "Benchmarking the accuracy of structure-based binding affinity predictors on Spike-ACE2 Deep Mutational Interaction Set"

**Supplementary Figures**

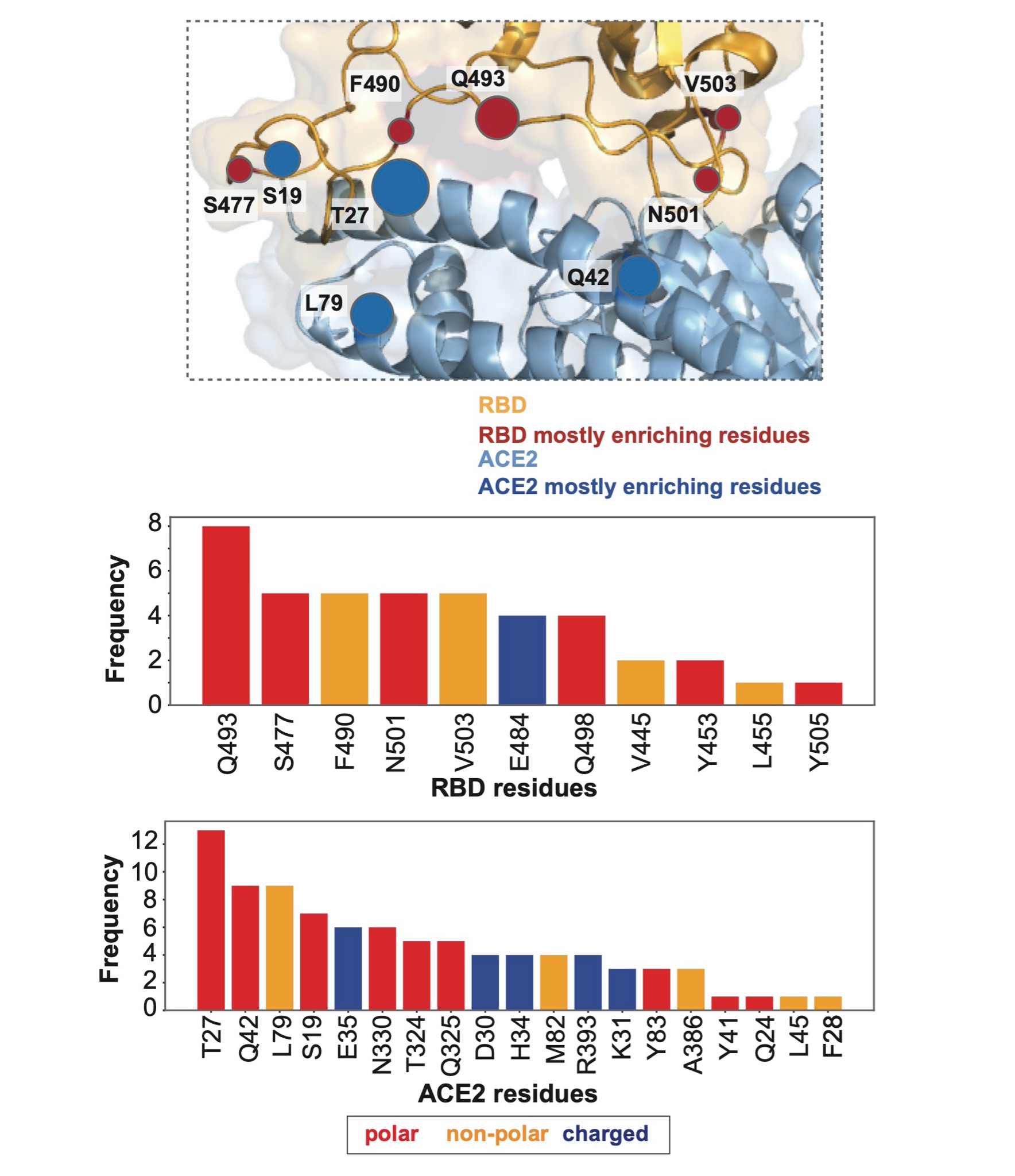

**Figure S1.** **The residues leading to most frequent binding enriching mutations.** In the bar plots, the bars are colored according to the residue’s physicochemical property (polar, non-polar and charged amino acids are shown in red, orange, and dark blue, respectively).

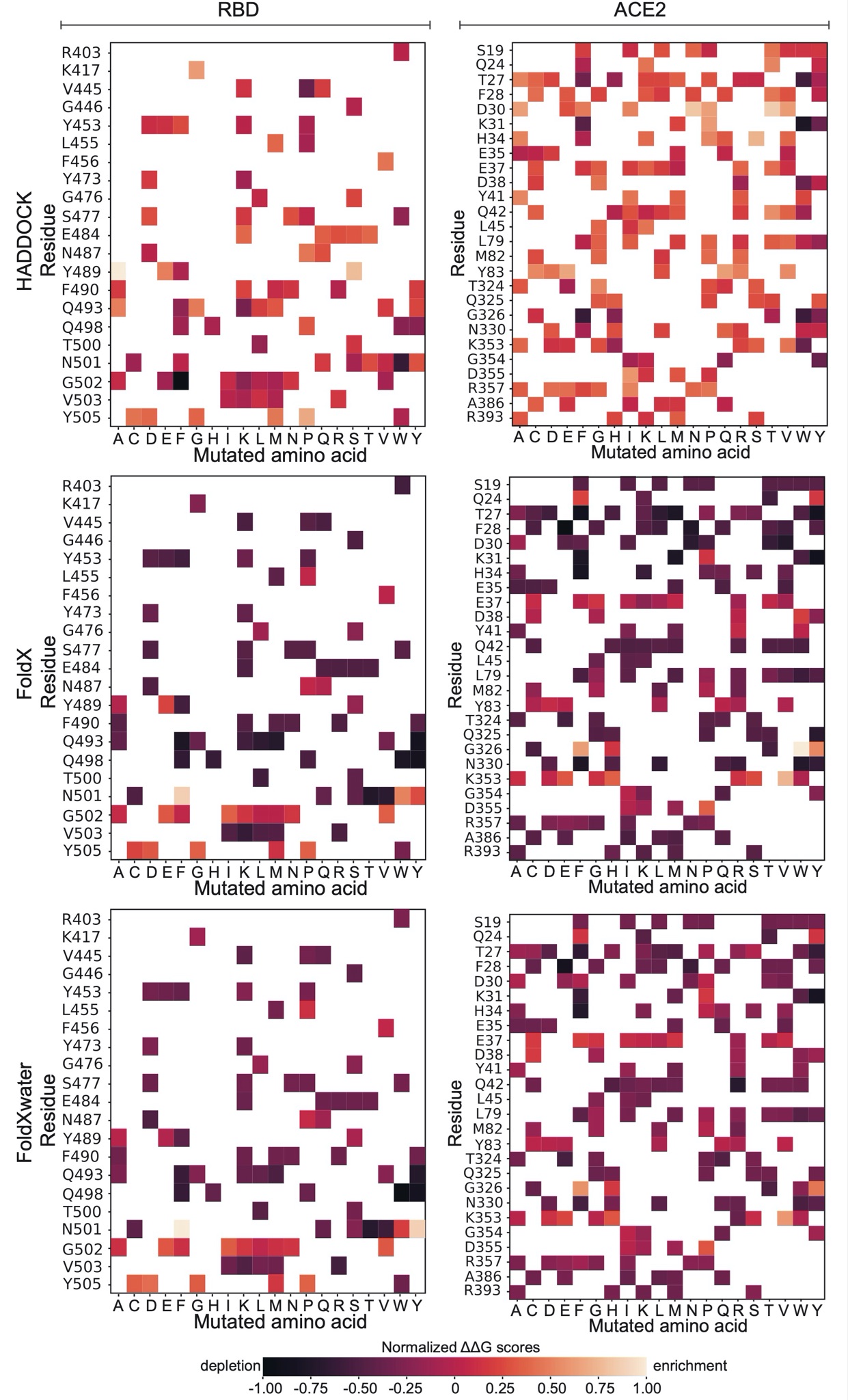

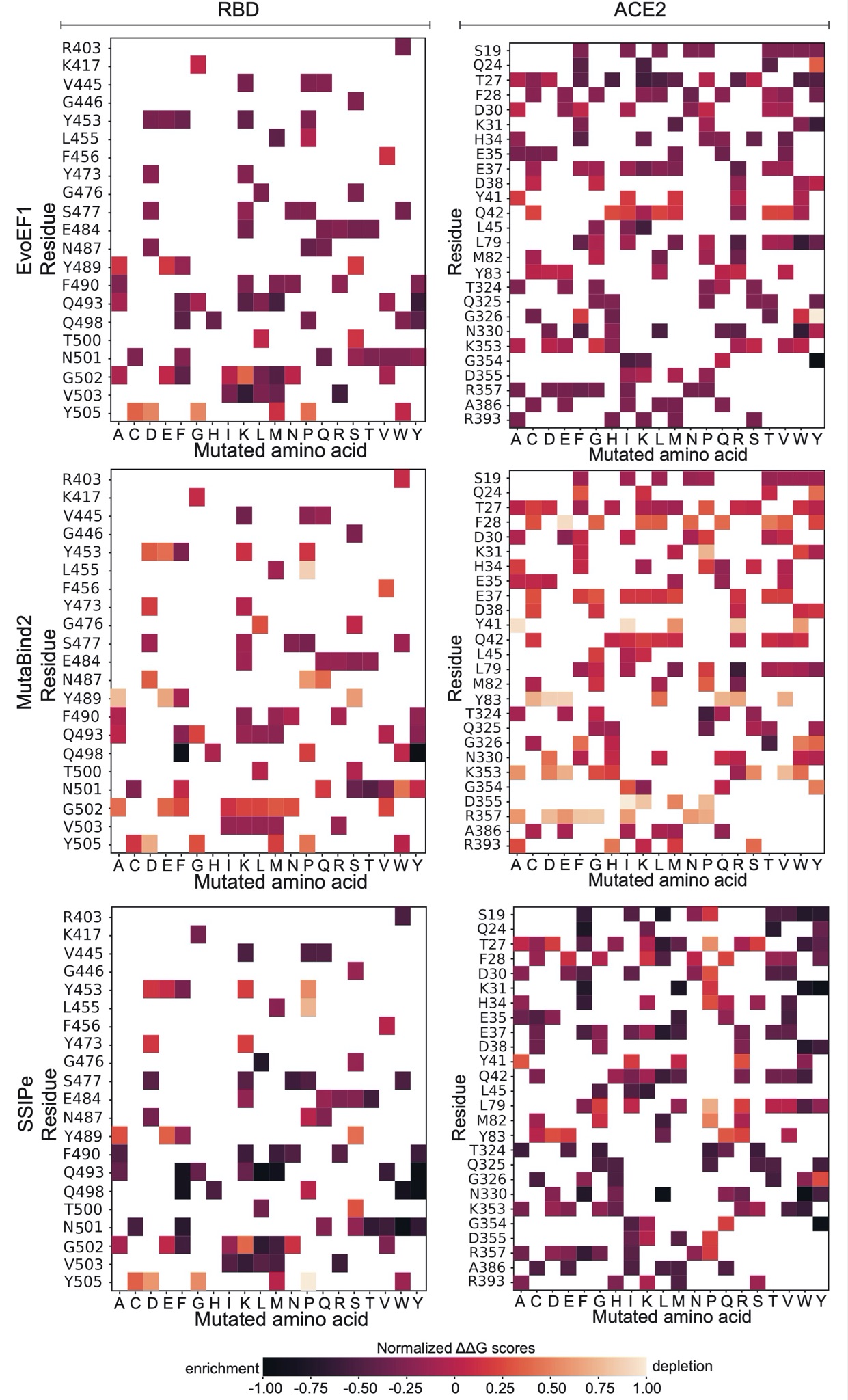

**Figure S2.** **Binding value change predictions** by HADDOCK, FoldX, FoldXwater, EvoEF1, MutaBind2, and SSIPe. The scores were normalized between 1 and -1 to use the same color scheme. The mutation is enriching if the predicted affinity change (∆Score) is <0 and depleting, if (∆Score) is >0. RBD Spike and ACE2 mutations are presented on the left and right columns, respectively.

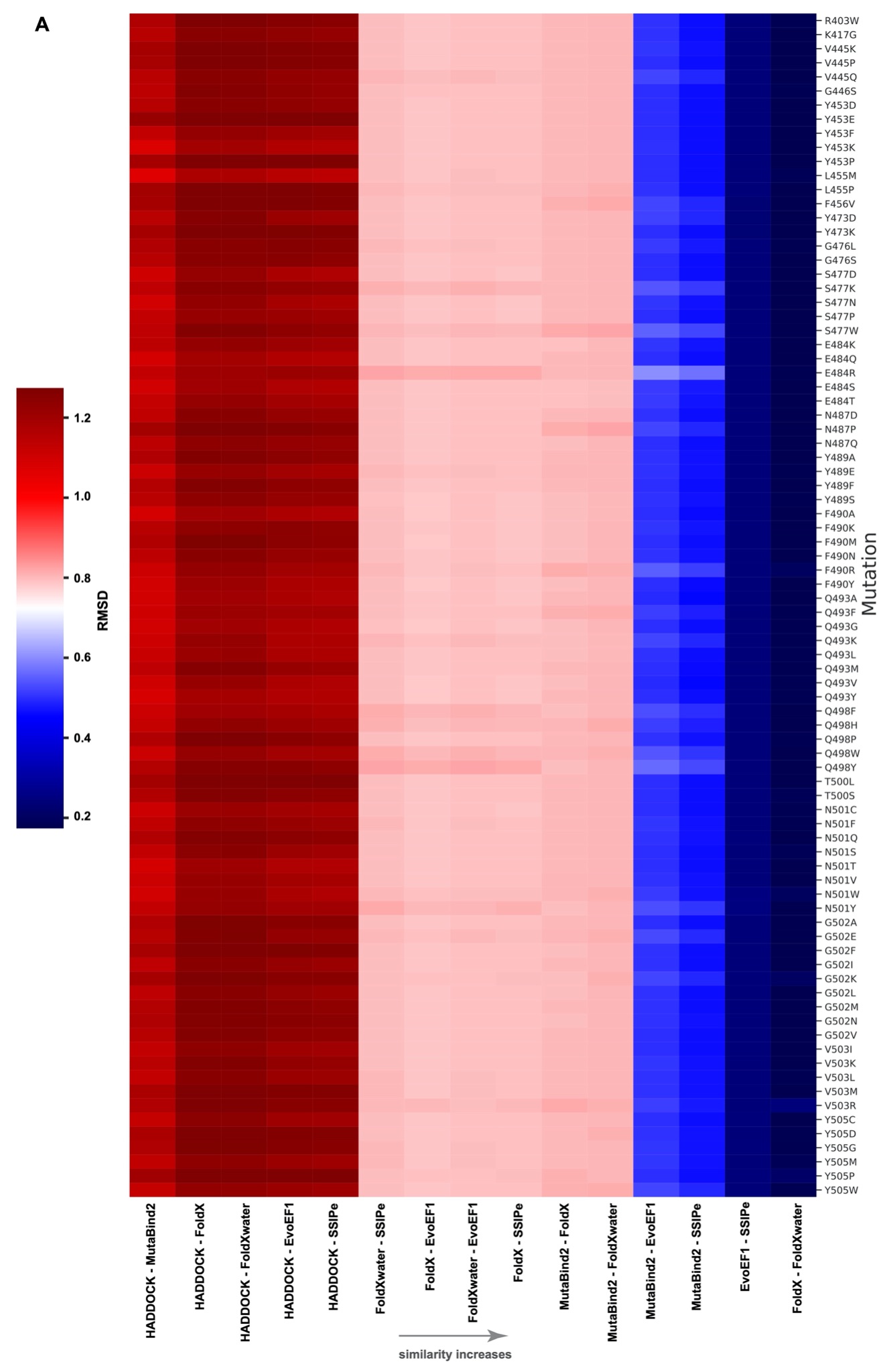

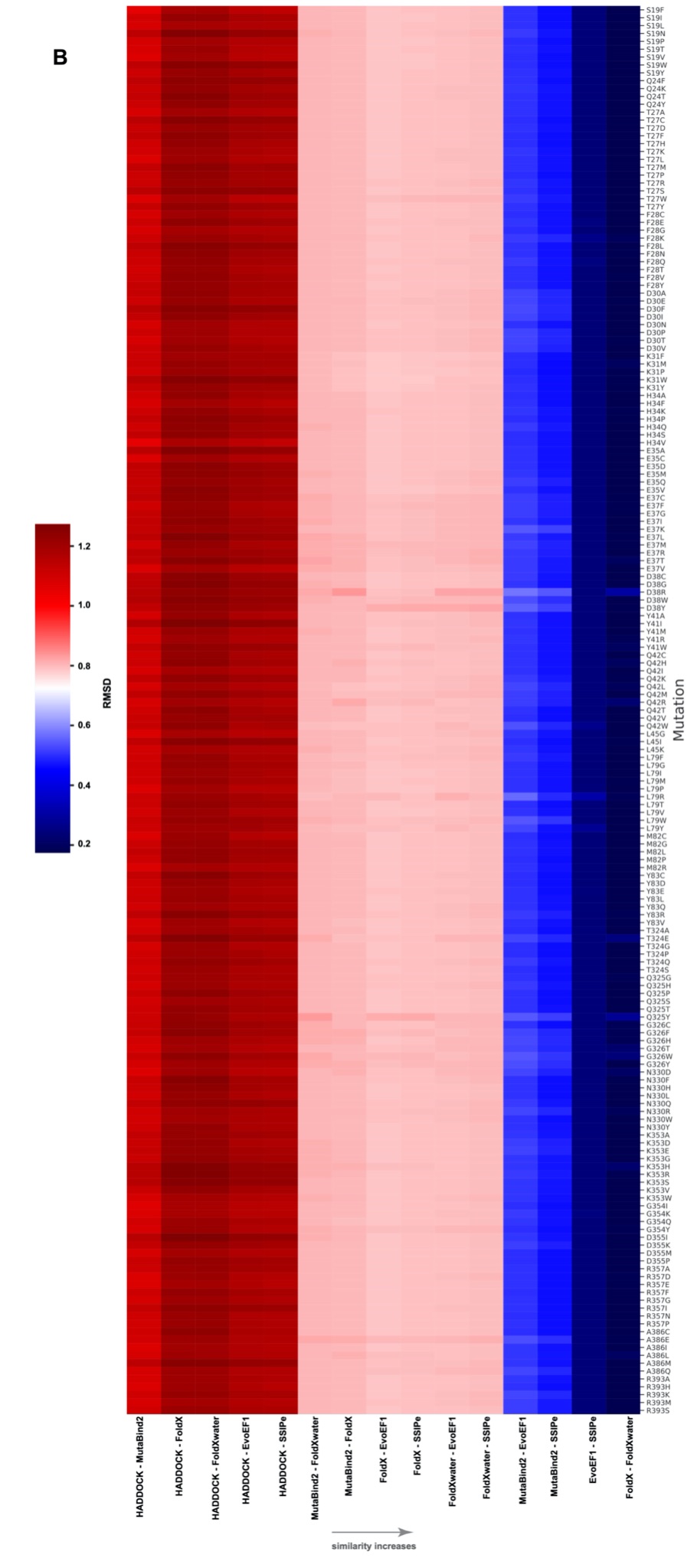

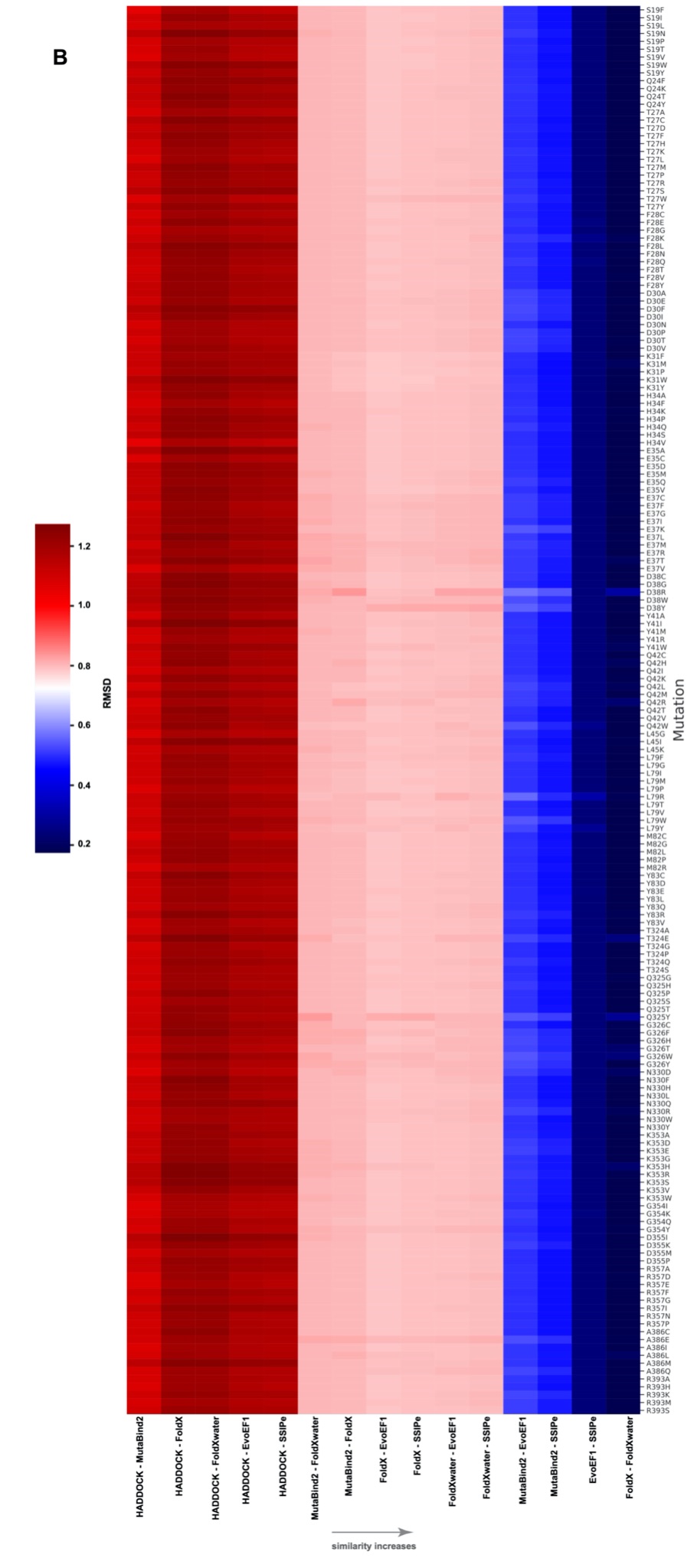
**Figure S3. Root Mean Square Deviations (RMSDs) of the variant complexes** for **(A)** RBD and **(B)** ACE2. RMSD values vary between 0 and 1.2 (Å) and the corresponding values are shown in blue and red, respectively.

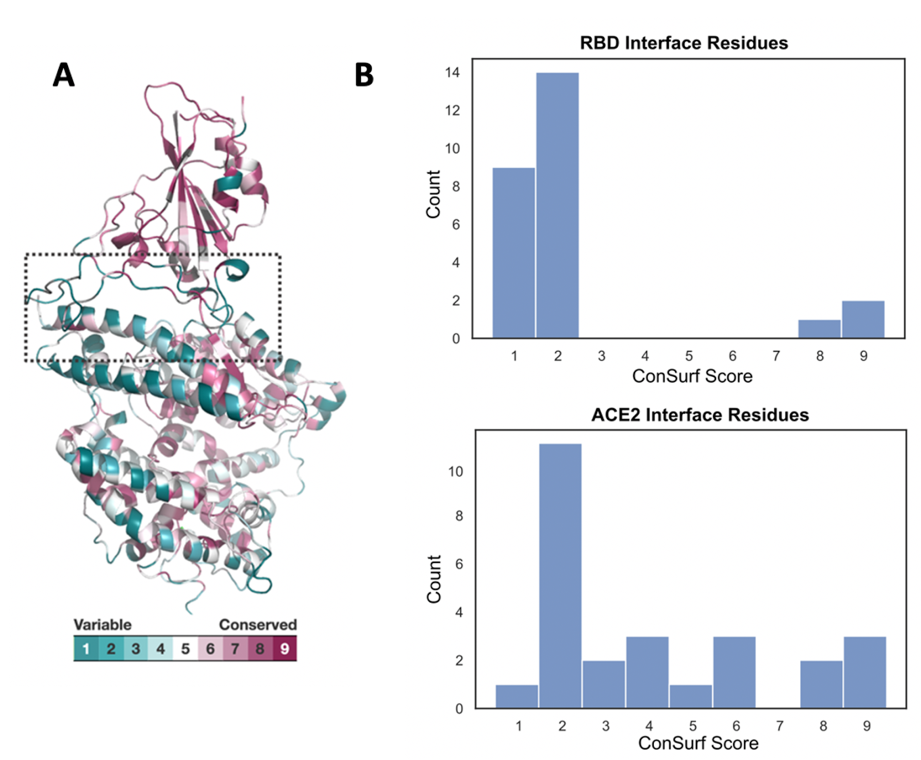

**Figure S4. Conservation analysis of the RBD-ACE2 complex (PDB ID: 6M0J) using ConSurf.** **(A)** Visual representation of the conservation profiles for RBD and ACE2 proteins. Conservation scores, ranging from 1 (indicating non-conserved residues) to 9 (indicating highly conserved residues), are color-coded in dark green and dark burgundy, respectively. **(B)** Bar plots displaying the conservation scores of interface residues for RBD (upper) and ACE2 (lower). The mean interface score is 2.3, which indicates a nonconserved interface.

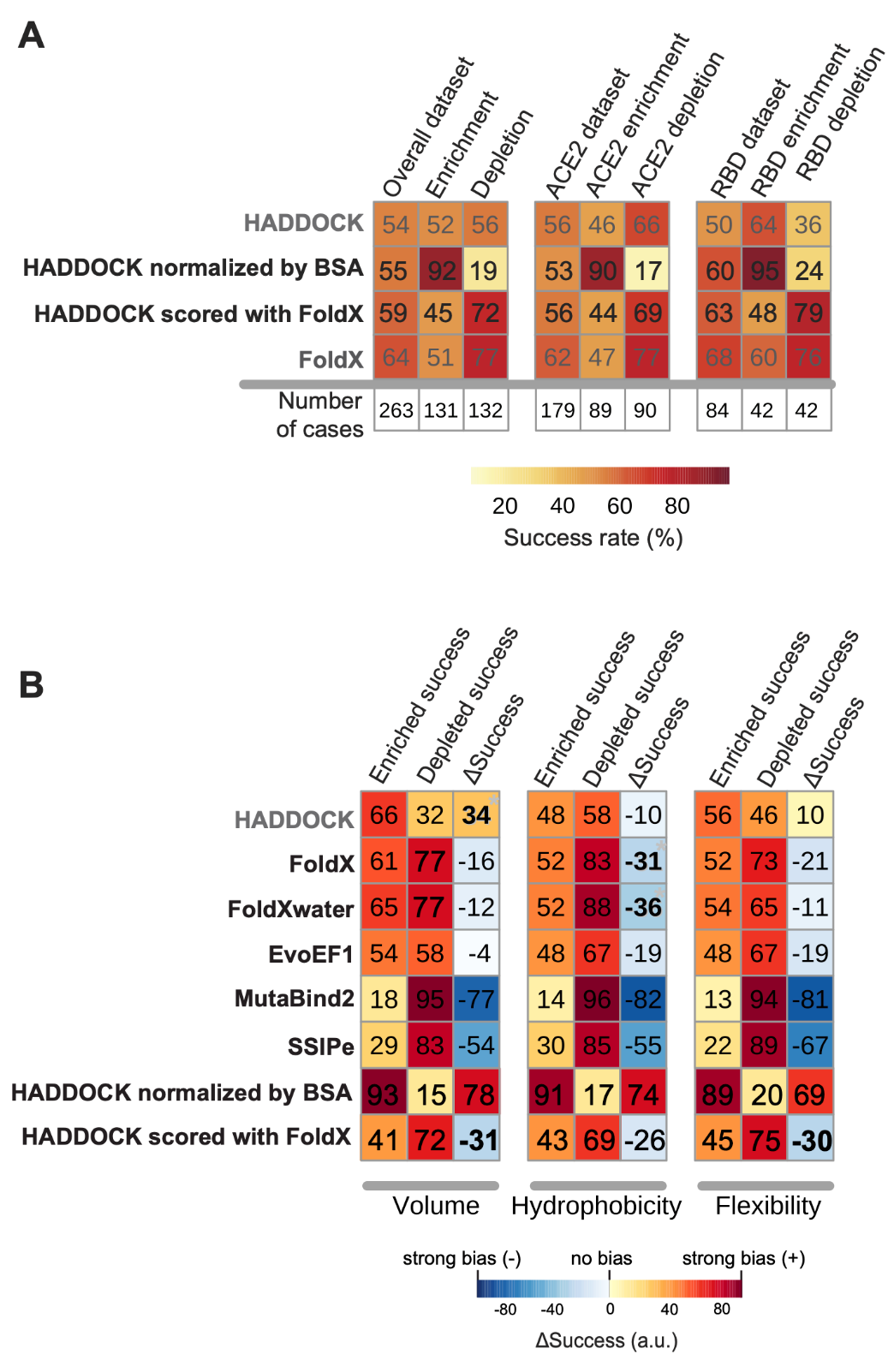

**Figure S5. The success rates** of HADDOCK, when its models are scored with HADDOCK scores normalized by Buried Surface Area (BSA) and with FoldX. The baseline FoldX and HADDOCK success rates are shown in gray.

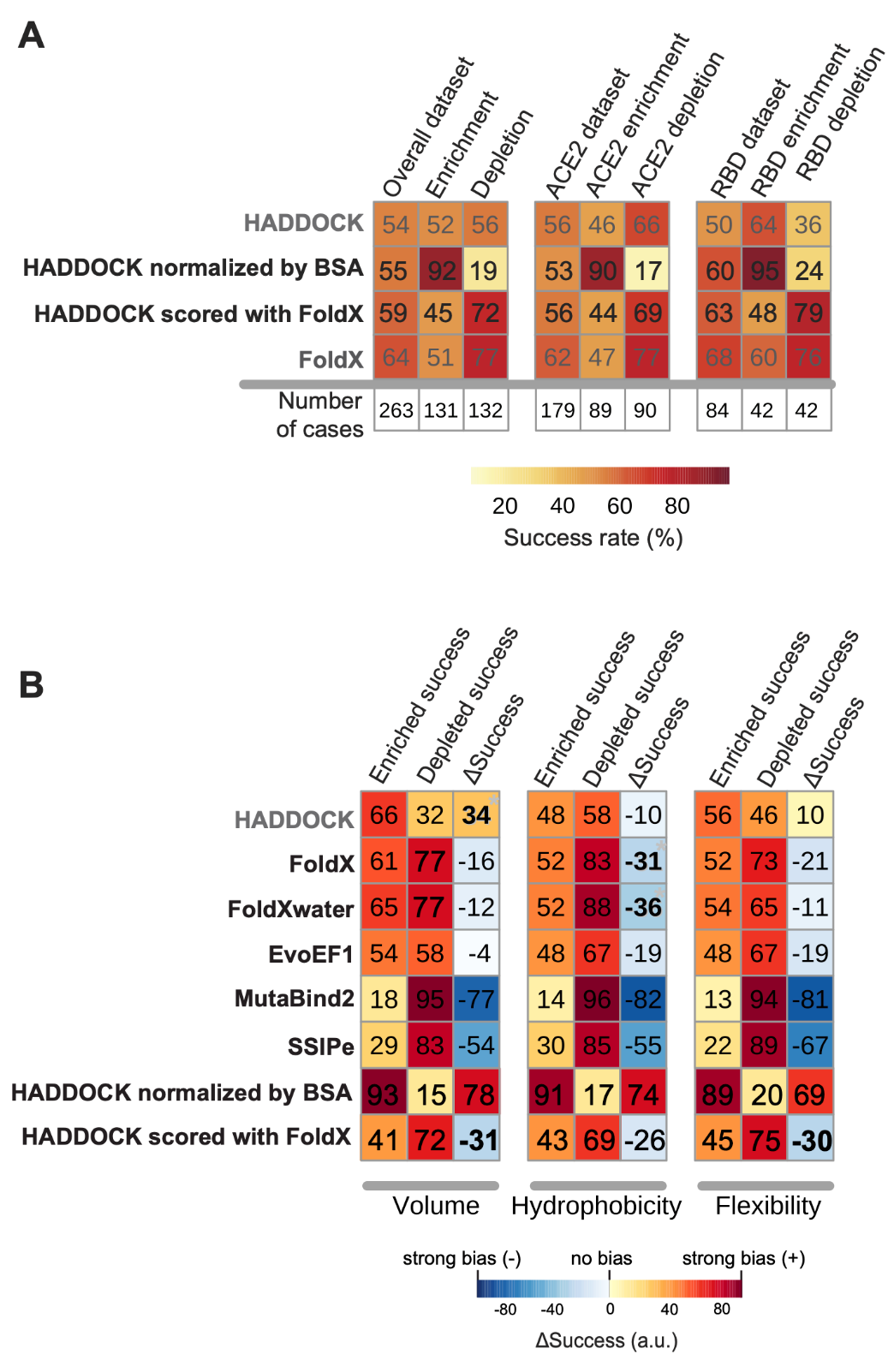

**Figure S6. The areas under the curve ratios** are calculated by using the frequency distributions plots given in Figure 4A-C. ∆Success rate is calculated by the change in the area under the curve for enriching and depleting class success curves (∆Success rate=Enriched Success – Depleted Success). Most extreme values are shown in bold.

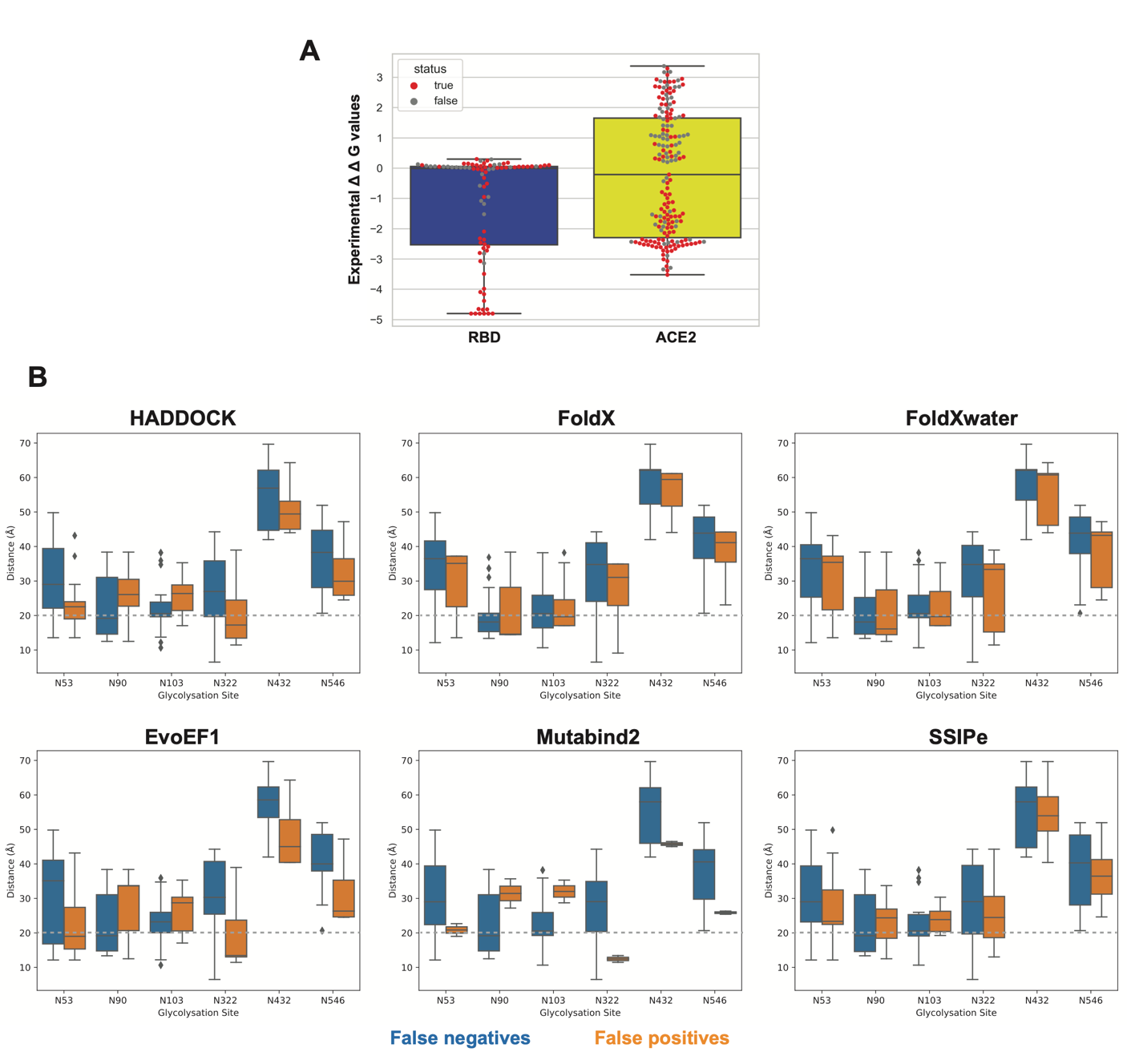

**Figure S7. (A) The distribution of correctly predicted FoldX data points on the experimental DMS dataset.** The correctly predicted cases are marked in red. **(B) The distance distributions of false negative (cases that were predicted as depleting even though they are enriching, blue) and false positive (cases that were predicted as enriching even though they are depleting orange) cases to the six ACE2 glycosylation sites (N53, N90, N103, N322, N432, and N546).** The distance distribution plots are presented for HADDOCK, FoldX, FoldXwater, EvoEF1, MutaBind2, and SSIPe predictors. A 20 Å threshold is denoted by a gray dashed line, representing the distance at which glycans can exert additional electrostatic effects.

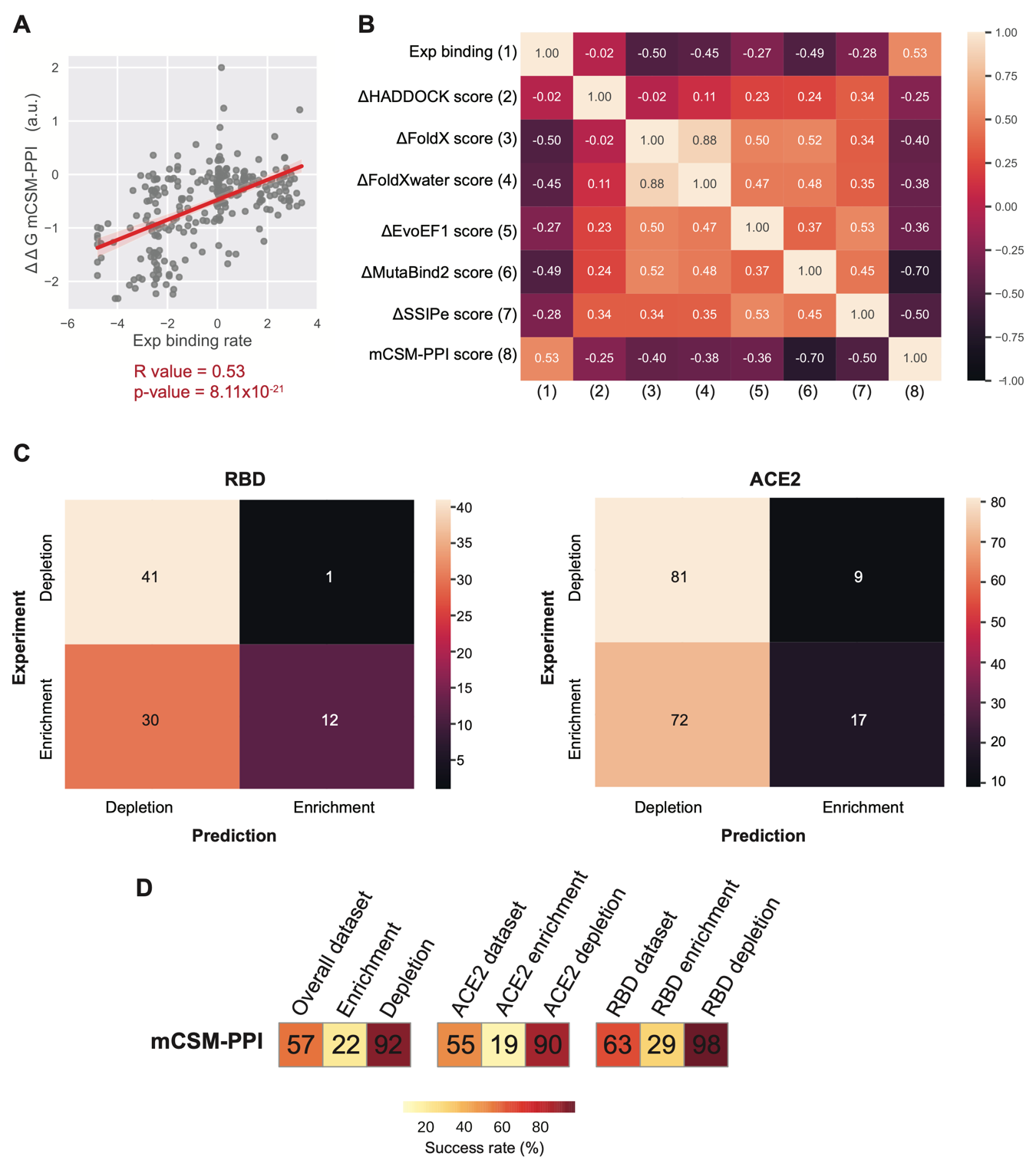

**Figure S8. mmCSM-PPI Results. (A)** Correlation between the experimental DMS benchmark set and mCSM-PPI results. The change in binding affinity is calculated as the wild-type (WT) value minus the mutant value. Therefore, a positive value indicates an increase in binding affinity, while a negative value signifies a decrease, consistent with the experimental data. That’s why the correlation is represented by R = 0.53, where p-value = 8.11x10-21. **(B)** Correlation heatmap depicting the relationship between computational and experimental scores. Pearson Correlation Coefficients (PCC) are used to measure the correlation, with values ranging from -1 to 1. **(C)** Confusion matrices for RBD (left) and ACE2 (right) predictions. The y-axis displays the DMS dataset results, while the x-axis shows the mmCSM-PPI2 prediction results. Darker colors signify unsuccessful predictions, while lighter colors indicate successful ones. **(D)** Success rates for the overall dataset, ACE2, and RBD datasets, respectively. Success rates range from 0 to 100, with lighter colors indicating successful results and darker colors indicating unsuccessful ones.

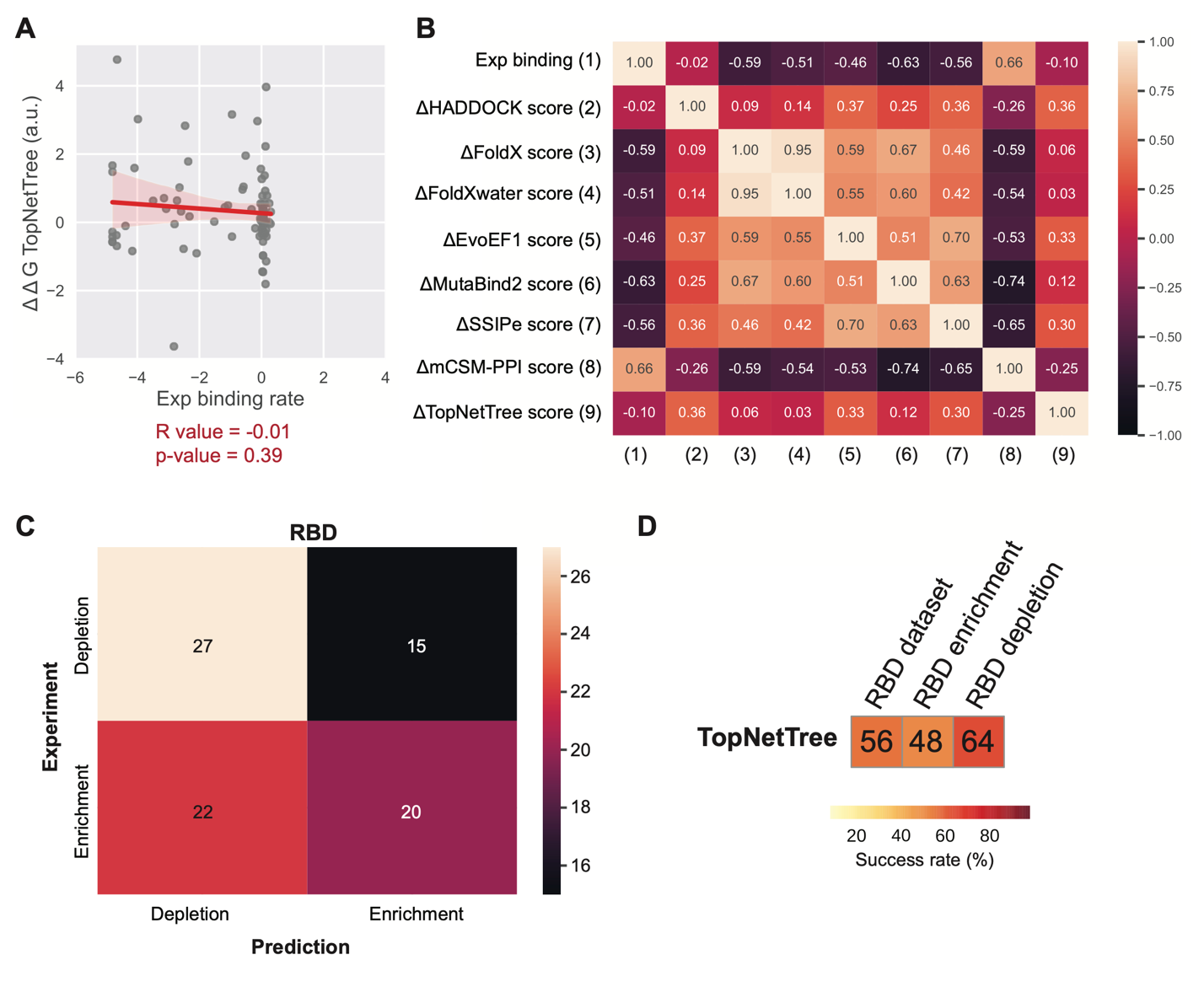

**Figure S9. TopNetTree Results. (A)** Correlation between the experimental RBD DMS benchmark set and TopNetTree predictions. The correlation is represented by R = -0.10 and p-value = 0.39. **(B)** Correlation heatmap depicting the relationship between computational and experimental scores for RBD dataset. Pearson Correlation Coefficients (PCC) are used to measure the correlation, with values ranging from -1 to 1. **(C)** Confusion matrix for RBD predictions. The y-axis displays the DMS dataset results, while the x-axis shows the TopNetTree prediction results. Darker colors signify unsuccessful predictions, while lighter colors indicate successful ones. **(D)** Success rates for the RBD datasets. Success rates range from 0 to 100, with lighter colors indicating successful results and darker colors indicating unsuccessful ones.

**Supplementary Tables**

**Table S1. The binding enriching substitutions and their experimental binding values. (A)** Experimental binding values of RBD. **(B)** Experimental binding values of ACE2. **The variants of concern are underlined.**

| **Q493** | **S477** | **F490** | **N501** | **V503** | **E484** | **Q498** |
| --- | --- | --- | --- | --- | --- | --- |
| **Q493M 0.18**  Q493A 0.13  Q493Y 0.12  Q493F 0.06  Q493V 0.05  Q493L 0.05  Q493K 0.05  Q493G 0.01 | **S477D 0.09**  S477P 0.06  S477N 0.06  S477K 0.03  S477W 0.02 | **F490K 0.09**  F490Y 0.06  F490A 0.03  F490N 0.02  F490R 0.01 | **N501F 0.29**  N501Y 0.24  N501V 0.15  N501W 0.11  N501T 0.10 | **V503M 0.10**  V503K 0.10  V503I 0.05  V503L 0.03  V503R 0.01 | **E484R 0.15**  E484K 0.06  E484T 0.05  E484Q 0.03 | **Q498H 0.30**  Q498Y 0.16  Q498F 0.15  Q498W 0.07 |

| **T27** | **Q42** | **S19** | **L79** |
| --- | --- | --- | --- |
| **T27L 2.85**  T27Y 2.70  T27M 2.55  T27C 2.42  T27H 2.34  T27F 2.29  T27A 2.11  T27W 2.11  T27D 2.10  T27K 1.65  T27S 0.77  T27P 0.51  T27R 0.25 | **Q42C 2.85**  Q42L 2.74  Q42M 2.48  Q42V 1.92  Q42K 1.73  Q42I 1.71  Q42H 1.40  Q42R 0.84  Q42T 0.39 | **S19P 2.33**  S19F 1.13  S19V 1.09  S19W 1.07  S19Y 1.05  S19I 0.77  S19T 0.72 | **L79T 3.37**  L79W 3.29  L79V 3.18  L79I 2.90  L79Y 2.86  L79F 2.68  L79P 1.86  L79R 1.06  L79M 1.03 |

**Table S2. Correlation values (R and p values) between the experimental DMS benchmark set and the binding affinity predictors.** The dataset comprises n=262 data points for all scenarios, except for UEP, which consists of 129 data points. Correlation plots corresponding to this data can be found in Figure 2A, S8A and S9A.

| **Predictor** | **R value** | **p-value** |
| --- | --- | --- |
| **HADDOCK** | -0.02 | 0.77 |
| **FoldX** | -0.51 | 3.30x10^-18^ |
| **FoldXwater** | -0.45 | 8.20 x10^-15^ |
| **EvoEF1** | -0.27 | 1.15 x10^-5^ |
| **MutaBind2** | -0.49 | 1.73 x10^-17^ |
| **SSIPe** | -0.28 | 2.88 x10^-6^ |
| **UEP** | 0.14 | 0.11 |
| **mCSM-PPI** | 0.53 | 8.11x10^-21^ |
| **TopNetTree** | -0.10 | 0.39 |

**Table S3.** The Van der Waals volume, hydrophobicity, and flexibility metrics used in this study.

| Amino Acid | Van Der Waals Volume  (nm^3^) | Hydrophobicity | Flexibility | Physicochemical Classification |
| --- | --- | --- | --- | --- |
| A | 0.05702 | 0.62 | 1 | Non-polar |
| R | 0.58946 | -2.53 | 81 | Charged |
| N | 0.22972 | -0.78 | 3 | Polar |
| D | 0.21051 | -0.90 | 3 | Charged |
| C | 0.14907 | 0.29 | 3 | Non-polar |
| Q | 0.34861 | -0.85 | 9 | Polar |
| E | 0.32837 | -0.74 | 9 | Charged |
| G | 0.00279 | 0.48 | 1 | Non-polar |
| H | 0.37694 | -0.40 | 3 | Charged |
| I | 0.37671 | 1.38 | 9 | Non-polar |
| L | 0.37876 | 1.06 | 9 | Non-polar |
| K | 0.45363 | -1.50 | 81 | Charged |
| M | 0.38872 | 0.64 | 27 | Non-polar |
| F | 0.55298 | 1.19 | 3 | Non-polar |
| P | 0.2279 | 0.12 | 2 | Non-polar |
| S | 0.09204 | -0.18 | 3 | Polar |
| T | 0.19341 | -0.05 | 3 | Polar |
| W | 0.79351 | 0.81 | 3 | Non-polar |
| Y | 0.6115 | 0.26 | 3 | Polar |
| V | 0.25674 | 1.08 | 3 | Non-polar |
